# Transcriptomic analysis reveals new reparative mechanisms of SCF+G-CSF-reduced neuropathology in aged APP/PS1 mice

**DOI:** 10.1101/2025.05.10.653288

**Authors:** Afua Addo, Bin Li, Sasidhar Murikinati, Robert Gardner, Li-Ru Zhao

**Affiliations:** Department of Neurosurgery, Norton College of Medicine, SUNY Upstate Medical University, Syracuse, New York, 13202, United States; Department of Neurology, Louisiana State University Health Sciences Center, Shreveport, Louisiana, 71103, United States

**Keywords:** Alzheimer’s disease, SCF+G-CSF, transcriptomic analysis, hematopoietic stem cells, S100a8, S100a9, cancer

## Abstract

Alzheimer’s Disease (AD) is a neurodegenerative disease characterized by amyloid plaque deposition, tau hyperphosphorylation, neuroinflammation, and cognitive decline. Our previous studies showed that combined treatment with stem cell factor (SCF) and granulocyte colony-stimulating factor (G-CSF) reduces AD pathology in APP/PS1 mice. This study aimed to explore the molecular mechanism underlying SCF+G-CSF’s therapeutic effects using transcriptomic analysis. Aged APP/PS1 mice received daily subcutaneous injections of SCF+G-CSF or vehicle for 12 days. RNA was extracted from brain tissue on day 13 for gene chip analysis. Age-matched wild-type (WT) mice served as controls. Data were analyzed using TAC, STRING v12.0, Reactome, and ShinyGO 0.77. A total of 45,037 differentially expressed genes (DEGs) were detected. Twenty-seven DEGs met a ≥2-fold threshold in SCF+G-CSF-treated vs. vehicle-treated APP/PS1 mice; 89 DEGs met this threshold in APP/PS1 vs. WT mice. SCF+G-CSF treatment upregulated six immune-related genes (S100a8, S100a9, Ngp, Lcn2, Ltf, and Camp) associated with amyloid clearance, immune cell recruitment, and repair. Pathway analysis showed downregulation of IL-2, IL-4, IL-7, and EGFR1, and upregulation of IL-17 signaling, suggesting modulation of both innate and adaptive immunity. Notably, SCF+G-CSF downregulated several oncogenes, including Cbl, Akap9, Kcnq1ot1, and Snhg11, highlighting an overlap between cancer and AD-related pathways. SCF+G-CSF also promoted NADPH oxidase activation via Rho GTPases and showed >400-fold enrichment in metal ion sequestration, indicating potential metal chelation effects. These findings suggest that SCF+G-CSF treatment modifies immune and metabolic pathways, reduces AD pathology, and highlights new therapeutic targets involving inflammation, metal homeostasis, and oncogenic signaling.

## 1. Introduction

Alzheimer’s Disease (AD) is a progressive neurodegenerative disorder characterized by extracellular amyloid beta (Aβ) plaque accumulation, intracellular tau hyperphosphorylation leading to neurofibrillary tangle (NFT) formation, activation of innate immune cells, neuroinflammation, and cognitive decline.^1,2^ Despite numerous efforts to target these hallmarks, AD remains the leading cause of dementia, projected to affect nearly 14 million Americans by 2060^3^, with costs of care exceeding $300 billion in 2020.^4^

Non-pharmacological interventions such as physical exercise and cognitive training have shown benefits in improving behavioral symptoms.^4^ However, the efficacy of current pharmacological treatments remains limited. FDA-approved cholinesterase inhibitors, NMDA receptor modulators, monoclonal antibodies targeting Aβ, NSAIDs, and cholesterol-lowering agents have largely failed to stop disease progression or demonstrate consistent clinical benefit in mouse models or AD patients.^2,4,5^

These outcomes underscore the need for new therapeutic approaches. Increasing attention has turned to the roles of innate and adaptive immune responses and neuroinflammation in AD pathogenesis. Microglia, the resident macrophages of the central nervous system (CNS), play a critical role in clearing Aβ and cellular debris and regulating neuroinflammatory signaling.^1,2,6^ Upon detecting pathological changes via pattern recognition receptors, microglia and other CNS immune cells such as astrocytes initiate inflammatory cascades involving cytokines like IL-1β, TNFα, and IL-6.^2^ While inflammation aids in clearing debris, its chronic activation can worsen disease progression.^2^

In healthy brains, Aβ is efficiently cleared by macrophages and innate immune cells, but this process becomes impaired with AD progression.^6^ Studies have also implicated cell death pathways, including apoptosis, pyroptosis, and necroptosis, in chronic inflammation, contributing to tau misfolding, impaired homeostasis, and reduced debris clearance.^2^ Thus, neuroinflammation is a core feature of AD and a frequent target of therapeutic strategies.

Beyond debris clearance, glial cells also mediate chemoattraction, recruitment, and migration of immune cells.^2^ This highlights the potential of modulating intrinsic glial functions and hematopoiesis, the process of generating blood and immune cells, ^7^ as a therapeutic avenue. Although hematopoietic system dysfunction is implicated in AD, ⁵ its therapeutic targeting remains underexplored. Hematopoietic growth factors such as stem cell factor (SCF) and granulocyte colony-stimulating factor (G-CSF) regulate the proliferation, migration, and differentiation of hematopoietic stem cells (HSCs).

Importantly, peripheral monocytes can cross the blood–brain barrier in aged APP/PS1 mice and selectively target amyloid plaques in the brain.^8^ Under inflammatory conditions, hematopoietic progenitor cells are activated in the bone marrow, differentiate into macrophages, and participate in amyloid clearance and brain repair. ^7,8^

Clinical studies reveal that rheumatoid arthritis patients exhibit a lower AD risk, potentially due to elevated circulating hematopoietic growth factors such as G-CSF, M-CSF, and GM-CSF levels.^5^ Recombinant GM-CSF (Sargramostim) has shown efficacy in AD patients by reducing amyloid burden, enhancing immune responses, and improving cognition.^5^

Peripherally derived macrophages also outperform microglia in amyloid clearance.^6^ Yan and colleagues observed an increased presence of peripherally derived macrophages in the choroid plexus, perivascular spaces, meninges, and around amyloid plaques in aged APP/PS1 mice.^8^ Clonal hematopoiesis of indeterminate potential (CHIP), an age-related HSC expansion, alters myeloid function and is associated with reduced AD risk.^6, 9^ Furthermore, levels of 18 hematopoietic modulators, including G-CSF and M-CSF, have been linked to the transition from mild cognitive impairment to AD.^10^

Our previous studies demonstrated that combined treatment with hematopoietic growth factors, SCF and G-CSF (i.e., SCF+G-CSF), significantly reduced Aβ burden, neuroinflammation, and enhanced synaptic and dendritic regeneration in middle-aged and aged APP/PS1 mice.^11,12^ SCF+G-CSF-treated aged APP/PS1 mice showed reduced compact and fibrillary Aβ deposits, lower pro-inflammatory markers such as NOS, increased anti-inflammatory IL-4, and elevated expression of TREM2, a microglial receptor essential for amyloid clearance. ^11^

Despite these promising results, the mechanisms underlying SCF+G-CSF treatment’s therapeutic effects remain unclear. Therefore, the objective of this study is to employ transcriptomic analysis to reveal the molecular mechanisms by which SCF+G-CSF mitigates AD pathology.

## 2. Materials and Methods

All animal procedures were conducted in accordance with the National Institute of Health Guide for the Care and Use of Laboratory Animals and were approved by the Institutional Animal Care and Use Committee.

### 2.1. Animals and Experimental Design

APPswe/PS1dE9 (APP/PS1) mice (stock# 34832) and C57BL/6J mice (Wild Type control mice) (stock# 000664) (The Jackson Laboratory) were used in this study. The APP/PS1 mice co-express Swedish double mutations (K595N/M596L) of amyloid precursor protein (APP) and mutant human presenilin protein 1 (PS1).

Our previous studies showed that 12-day treatment with SCF+G-CSF (s.c.) mitigated AD neuropathology in both middle-aged and aged APP/PS1 mice ^11,12^; therefore, this study followed the same treatment paradigm to explore the mechanism of SCF+G-CSF treatment in aged APP/PS1 mice with advanced AD neuropathology using gene chip technology.

APP/PS1 mice (male, 17-18 months old) were randomly divided into two groups: a vehicle control group and an SCF+G-CSF treatment group (n=3). Age and sex-matched C57BL/6J mice served as wild-type (WT) controls (n=2). Recombinant mouse SCF (200μg/kg/day; PeproTech, USA) and recombinant human G-CSF (50μg/kg/day; Amgen, USA) or an equal volume of saline were subcutaneously (s.c.) injected for 12 consecutive days. One day (day 13) after the final injection, mice were euthanized. Brain tissue from the frontal lobes was dissected for RNA extraction and used for gene chip analysis. Following dissection, brain tissue was rapidly frozen on dry ice and stored at −80°C until further processing.

### 2.2. RNA Extraction Method

Frozen brain tissue was transferred into RNA*later* stabilization solution (Catalog# AM7021, ThermoFisher, USA) at a ratio of 100 mg of tissue per 500 μl of RNA*later*. RNA extraction was performed using the RNeasy Lipid Tissue Mini Kit (Catalog# 74804, Qiagen, USA). Briefly, after removing the brain tissue from RNA*later*, it was homogenized in 1 ml of QIAzolL Lysis Reagent using a glass homogenizer (≤ 100 mg brain tissue per sample).

The homogenate was incubated at room temperature (15-25°C) for 5 min, then centrifuged at 14,000 x g for 30 seconds. The resulting homogenate was transferred to a MaXtract tube along with 70μl RNase-free water. Next, 200μl of Chloroform was added, the tube was shaken vigorously for 15 seconds, incubated at room temperature for 2 – 3 min, and centrifuged at 12,000 x g for 15 min at 4°C.

The upper, aqueous phase was transferred to a new tube, mixed with 600μl of 70% ethanol, and vortexed. The sample was then loaded onto an RNeasy Mimi spin column placed in a 2 ml collection tube and centrifuged at 8,000 x g for 15 seconds at room temperature. The flow-through was discarded.

To wash the membrane, 350μl Buffer RW1 was added to the spin column and centrifuged at 8,000 x g for 15 seconds. The flow-through was discarded. A DNase digestion step was performed by mixing 10μl of DNase I stock solution with 70μl of Buffer RDD (Cat# 79254, RNase-Free DNase Set, Qiagen, USA). After gentle inversion, 80μl of the DNase 1 mix was applied directly to the spin column membrane and incubated for 15 min at room temperature.

Following DNase digestion, 350μl of Buffer RW1 was added to the spin column, centrifuged at 8,000 x g for 15 seconds, and the flow-through was discarded. The membrane was then washed with 500μl of Buffer RPE, centrifuged at 8,000 x g for 15 seconds, and the flow-through was discarded. A second wash was performed using the same volume of Buffer RPE, followed by centrifugation at 8,000 x g for 2 min.

The spin column was transferred to a new 2 ml collection tube and centrifuged at full speed for 1 min to remove residual buffer. It was then placed in a new 1.5 ml collection tube. To elute RNA, 30μl of RNase-free water was added directly to the column membrane and centrifuged at 8,000 x g for 1 min.

RNA integrity and purity were assessed using an Agilent Bioanalyzer before proceeding with gene chip analysis.

### 2.3. Gene Chip Data Analysis

Transcriptomic analysis of the gene chip data was performed using ThermoFisher Scientific’s Transcriptome Analysis Console (TAC).^13^ “.cel” files containing the microarray data for mice labelled APP1, APP2, APP3, CON1, CON2, CON3, WT1, and WT2 were uploaded into TAC for gene expression comparison analysis after selecting the Mouse 430_2.0 array type, with the genome version set to “Mm10 (Mus musculus)”. Two separate analyses were performed: (1) SCF+G-CSF-treated APP/PS1 mice (APP) vs. vehicle-treated APP/PS1control mice (CON) and (2) CON vs WT controls. The analysis type was set to “Expression (gene)”, and RMA was selected for summarization. The one-way ANOVA empirical Bayes (“ebayes”) method was used for data analysis, and a 2.0-fold expression threshold was applied with a gene-level p-value cutoff of < 0.05. Out of the 45,037 genes detected from the microarray “. cel” files, 27 genes and 89 genes were significantly differentially expressed in the APP vs. CON and CON vs. WT comparisons, respectively.

The resulting differently expressed gene lists were entered into publicly available gene analysis tools including STRING v12.0^14^, ShinyGO v0.77^15^ (developed by South Dakota State University), and Reactome^16^ for further analysis.

## 3. Results and Discussion

Given that our previous studies demonstrated that SCF+G-CSF treatment mitigates AD neuropathology in the aged APP/PS1 mice ^11,12^, we aimed to explore the underlying mechanism of SCF+G-CSF treatment in this model at an advanced stage of disease using transcriptomic analysis. To accomplish this, 17–18-month-old APP/PS1 mice received subcutaneous (s.c.) injections of SCF+G-CSF or the vehicle solution for 12 consecutive days. One day after the final injection, RNA was extracted from the frontal lobes and used for gene chips analysis (Figure 1).

**Figure 1.**
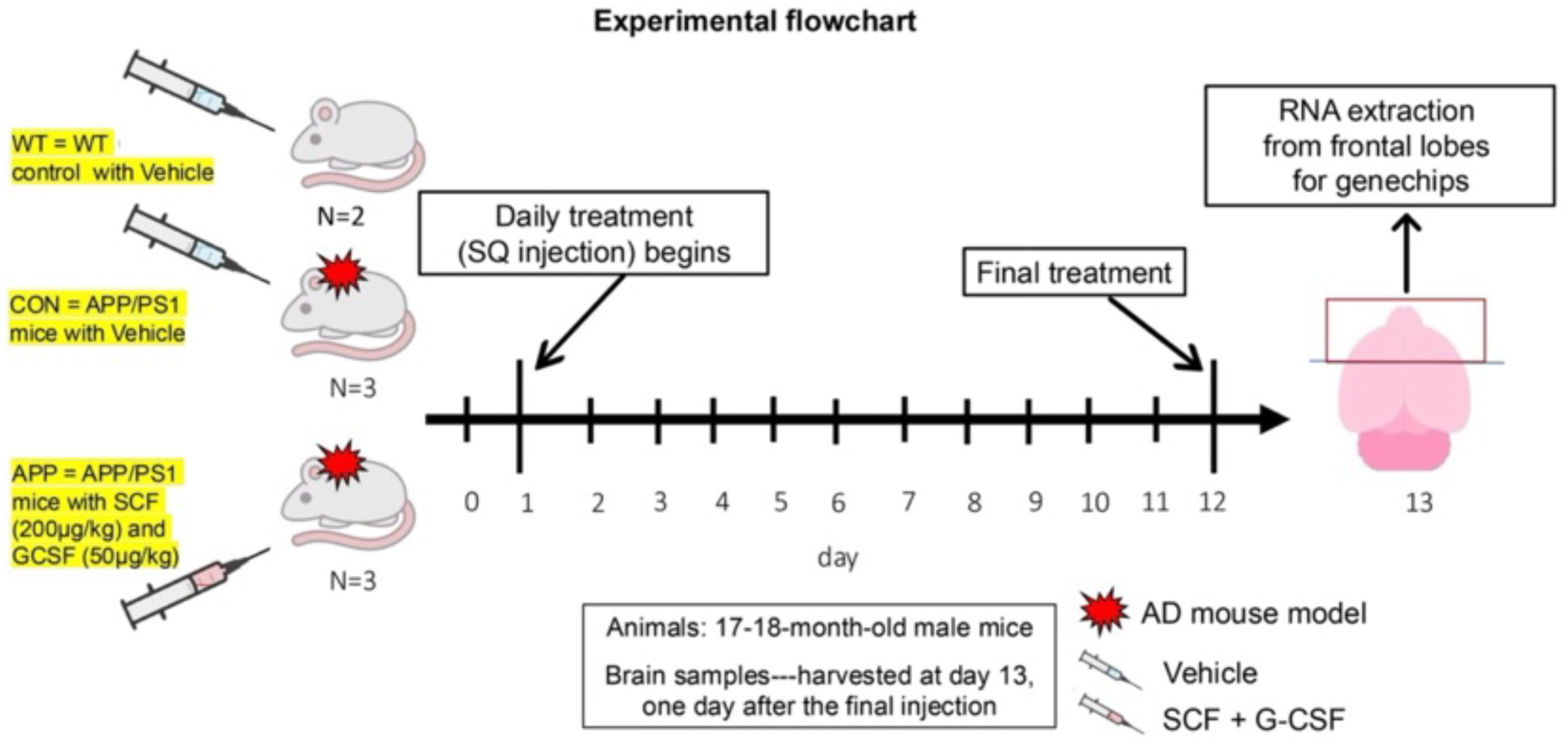
Schematic of experimental design. Male APP/PS1 mice (17-18 months old) were randomly divided into two groups (n=3/group): one received SCF+G-CSF treatment and the other received vehicle injections. Age-matched WT mice (n=2) receiving vehicle injections served as controls. All mice received daily subcutaneous (SQ) injections of SCF+G-CSF or vehicle for 12 consecutive days. On day 13, mice were euthanized, and RNA was extracted from frontal cortices for transcriptomic analysis using gene chip microarrays.

The APP/PS1 mouse model is widely used to study AD. This doubly transgenic model carries mutations in both the human APP and PS1 genes, leading to age-dependent amyloid plaque formation. In this model, plaque deposition and gliosis typically begin as early as 6 weeks of age, with cognitive impairment emerging around 7 months.^17^ For this present study, we used even older APP/PS1 mice to assess the transcriptomic profile of SCF+G-CSF treatment at a more advanced or chronic stage of AD, which we believe more accurately reflects the AD patient population and thus provides a more relevant evaluation of SCF+G-CSF’s therapeutic efficacy.

### 3.1. SCF+G-CSF treatment suppresses the transcriptional progression of Alzheimer’s disease in APP/PS1 mice, acting through the regulation of key genes

Gene expression analysis using ThermoFisher Scientific’s TAC platform revealed that SCF+G-CSF treatment induces differential expression in a small set of genes (Figure 2A). A list of differentially expressed genes (DEGs) in SCF+G-CSF-treated APP/PS1 mice (APP) compared to untreated APP/PS1 mice (CON) is presented in Table 1. Out of the 45,037 genes identified, only 27 genes were significantly differentially expressed and met the two-fold expression change threshold. Among these 27 DEGs, 19 were downregulated and 8 were upregulated following SCF+G-CSF treatment (Figure 2A, 2B). A heatmap representing hierarchical clustering of the dataset is shown in Figure 3, with red indicating upregulated genes and blue indicating downregulated genes.

**Figure 2.**
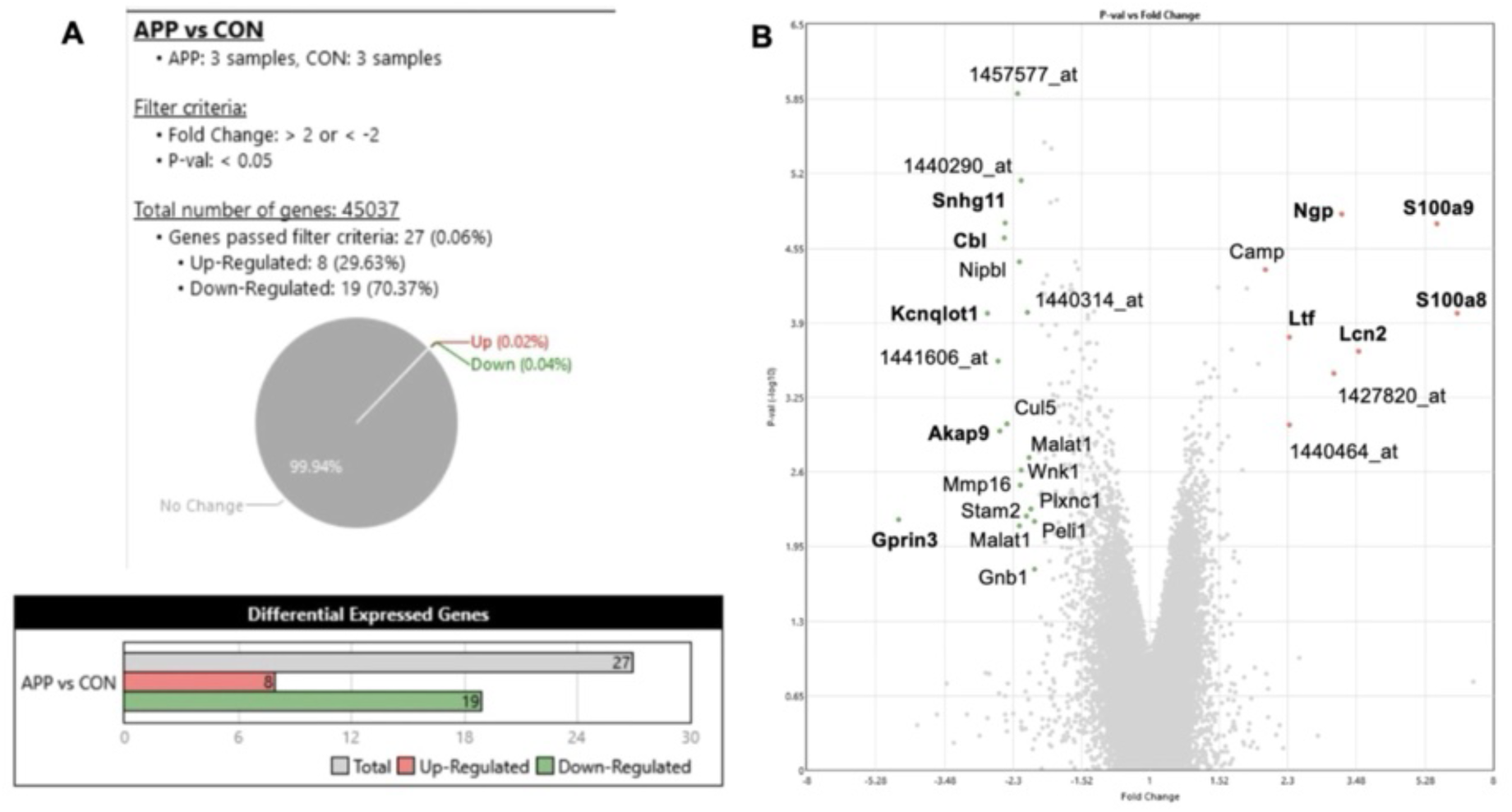
Summary and volcano plot of gene chip analysis using Transcriptome Analysis Console (TAC) in aged APP/PS1 mice treated with or without SCF+G-CSF. (A) SCF+G-CSF treatment significantly altered the expression of 27 genes, with over 70% being downregulated. (B) The top 5 upregulated and downregulated differentially expressed genes (DEGs) are highlighted in bold. Genes labelled with numerical probe set IDs lack known gene names or symbols. APP: APP/PS1 mice treated with SCF+G-CSF. CON: APP/PS1 control mice receiving vehicle injections.

**Table 1.**
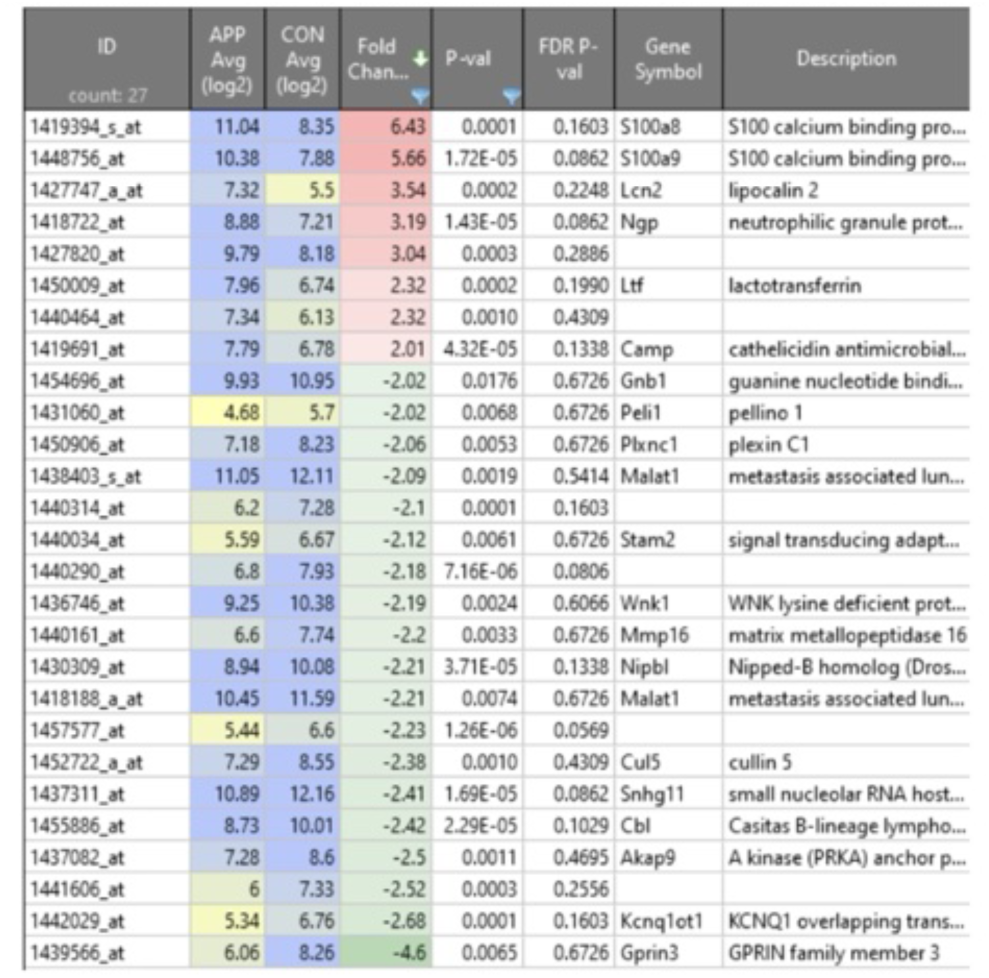
SCF+G-CSF-induced DEGs in aged APP/PS1 mice.

**Figure 3.**
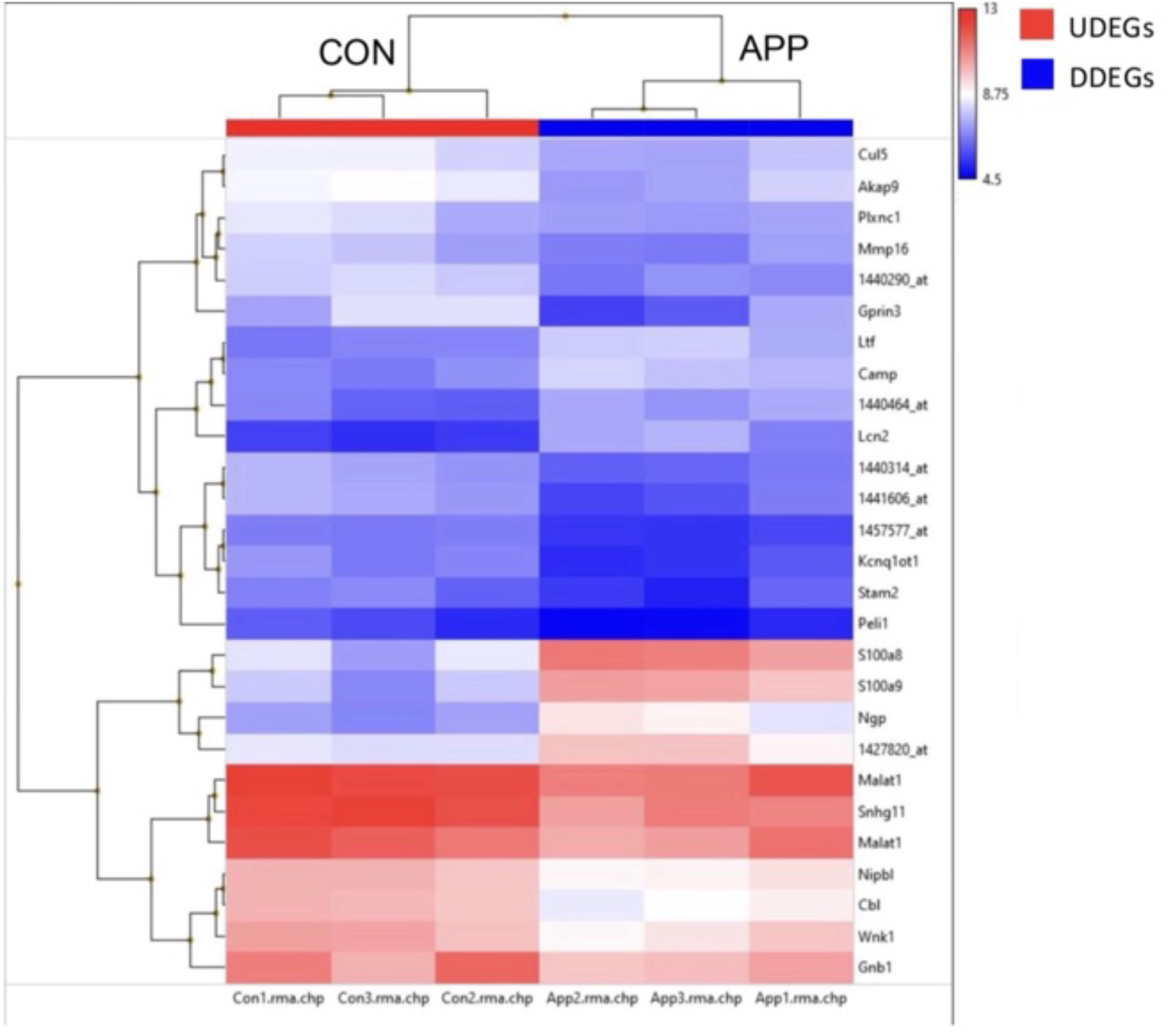
Heat map showing hierarchical clustering of differentially expressed genes DEGs in the brains of aged APP/PS1 mice treated with or without SCF+G-CSF. Normalized expression levels of DEGs are presented on a log-scale gradient, with darker colors indicating higher expression and lighter colors indicating lower expression. Red denotes upregulated DEGs (UDEGs), and blue indicates downregulated DEGs (DDEGs). The heat map was generated using the Transcriptome Analysis Console (TAC). APP: APP/PS1 mice treated with SCF+G-CSF. CON: APP/PS1 control mice receiving vehicle injections.

Analysis using ShinyGO v0.77 revealed that significantly more coding genes were differentially expressed in the APP/PS1-SCF+G-CSF (APP) group compared to APP/PS1 controls (CON) than would be expected by chance (p = 0.0025, Figure S1A). Coding genes provide the genetic instructions that are transcribed into mRNA and then translated into proteins.^18^ Our dataset also showed that a few genes related to long non-coding RNAs (IncRNAs) were significantly altered following SCF+G-CSF treatment (Figure S1A). IncRNAs play powerful roles in transcriptional regulation, chromatin modification, and post-translational control.^19^ These findings suggest that SCF+G-CSF treatment preferentially modulates the expression of coding genes while also impacting lncRNAs, thereby influencing both transcriptional and translational processes and altering cellular functions. Figure S1B illustrates a chromosomal map of DEGs identified from TAC analysis, further confirming that SCF+G-CSF treatment predominantly drives gene downregulation. The base pair localization of each gene on its respective chromosome is provided in Table S1. Together, these data indicate that most genes affected by SCF+G-CSF treatment are suppressed coding genes in aged APP/PS1 mice. Thus, even a modest transcriptional shift or “transcriptional hold” may be sufficient to reduce AD-related neuropathology.

### 3.2. Functions and Implications for SCF+G-CSF–Upregulated Differentially Expressed Genes (UDEGs)

All UDEGS are shown in Figure 2B and Table 1. The functions and implications of the 6 known UDEGs following SCF+G-CSF treatment are inferred from transcriptomic analysis of gene chip data and supported by findings from previous studies.

#### S100a8 and S100a9

The two most UDEGs following SCF+G-CSF treatment in our dataset are *S100a8* and *S100a9*, which are upregulated 6.43-fold and 5.66-fold, respectively (Table 1). These genes encode cytoplasmic calcium-binding proteins that can form either homodimers or heterodimers with each another. Once secreted into the extracellular space, S100A8 and S100A9 can bind to amyloid.^20,21^

It has been reported that S100A8/A9 proteins are associated with proinflammatory states and promote amyloid formation.^22^ However, our findings, along with previous studies, do not fully support this view. The localization of S100 proteins to amyloid plaques does not necessarily indicate a direct role in amyloidogenesis, as the functions of this protein family are far more diverse. Firstly, S100A8 and S100A9 proteins are critically involved in the recruitment of arachidonic acid from the cell membrane.^20,23^ They bind to the NADPH oxidase (NOX) enzyme complex and contribute to the oxidative burst required for the clearance of invading organisms and the reduction of oxidative stress.^20,23^ Additionally, accumulating evidence suggests that upregulation of S100A9 promotes the differentiation of promyelocytes into myelocytes or granulocytes, facilitating myeloid maturation.^20^ This process is critical for the CNS response to amyloid deposition, as bone marrow- or myeloid-derived cells, such as macrophages, can adopt microglia-like phenotypes. These cells act as CNS phagocytes and are key contributors to amyloid clearance.^24^

Furthermore, S100A8 and S100A9 chelate Zn^2+^ to inhibit bacterial growth and also interact with microtubules to promote phagocyte migration.^21^ A study using *S100a9* knockout mice demonstrated reduced S100A8/A9 complex formation, along with impaired phagocytotic migration and delayed wound healing.^21^ Another study reported that S100A9 may act through a microtubule-associated pathway to prevent excessive phagocytosis in leukocytes.^25^ This mechanism resembles the microtubule-related processes involved in neurofibrillary tangle (NFT) formation, which are critical for intracellular trafficking. Collectively, these functions support our findings that SCF+G-CSF treatment promotes enrichment of genes involved in antimicrobial activity and metal ion binding.

Our study reveals that SCF+G-CSF-induced upregulation of *S100a8/a9* may contribute to proinflammatory states that activate and facilitate myeloid cell migration to amyloid plaque sites. We postulate that SCF+G-CSF-mediated enrichment of the IL-17 signaling pathway, another key proinflammatory cascade, may enhance the activity of S100A8/A9, thereby increasing the infiltration of innate and adaptive immune cells into the brain in response to AD-related pathology in aged APP/PS1 mice (Figure S2).

These findings align with previous reports demonstrating that microglia are both activated by and secrete inflammatory cytokines such as IL-6 and TNFα in a TLR-4-dependent manner.^26^ Notably, another study showed that these cytokines are required for the induction and formation of the neuroprotective and regenerative M2 microglial phenotype, which is known for promoting healing and repair in the CNS.^27^

The diverse functions of S100A8 and S100A9, as demonstrated by our findings and previous studies, suggest that these proteins are more likely involved in processes that target amyloid, recruit immune cells, and facilitate amyloid clearance, rather than in amyloid production.^20,21,23^ We propose that these SCF+G-CSF-upregulated genes play a critical role in maintaining a proinflammatory state necessary for microglial activation, migration, and localization to sites of amyloid deposition, working in concert with other genes in the SCF+G-CSF-upregulated cluster (Figure 4). SCF+G-CSF-mediated upregulation of *S100a8* and *S100a9* genes likely contributes to mitigating extracellular drivers of AD neuropathology by stimulating the immune system, promoting differentiation of immune cells, directing immune cells to amyloid plaques, enhancing amyloid degradation, and activating antimicrobial defenses in aged APP/PS1 mice. Although previous studies have reported increased S100A9 expression around amyloid plaques,^28^ we argue that the proximity of S100A8/A9 proteins to amyloid does not imply a causal role in amyloid production or stabilization.

**Figure 4.**
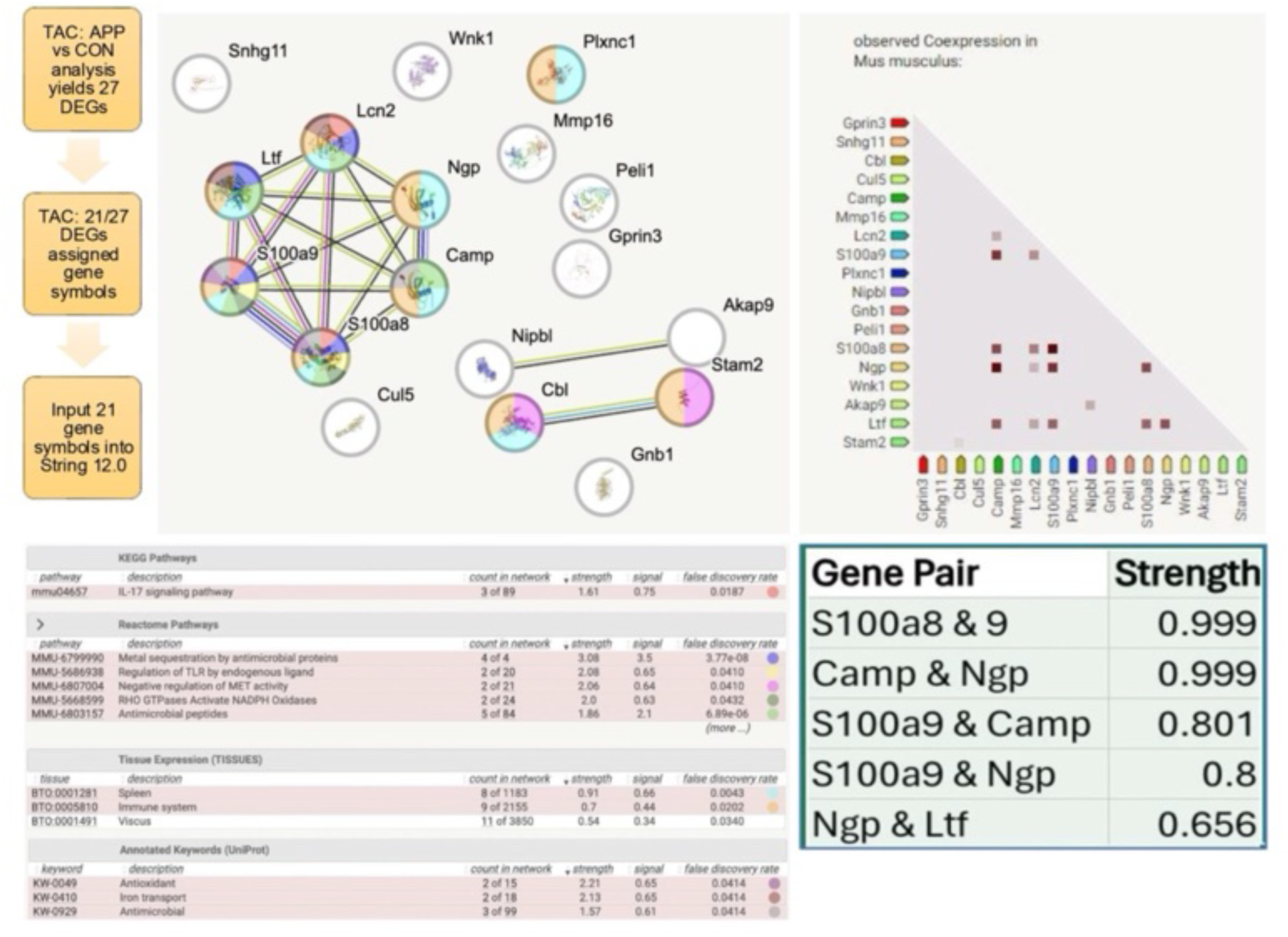
STRING 12.0 analysis of SCF+G-CSF-induced differentially expressed genes (DEGs) in the brains of aged APP/PS1 mice. Transcriptome Analysis Console (TAC) analysis identified 27 DEGs in response to SCF+G-CSF treatment, and 21 had identifiable gene names. Of these 21 DEGs, 18 were identified in the STRING 12.0 interaction analysis. SCF+G-CSF promoted both interaction and co-expression within a six-member gene cluster previously associated with spleen and immune tissue, now identified in the brains of APP/PS1 mice, suggesting that immune system dysregulation contributes to AD neuropathology. The identified genes are involved in pro-inflammatory signaling linked to microglia activation and phagocytosis, Aβ and neurofibrillary tangle recognition and localization, metal ion binding, and myeloid-mediated wound healing and cell growth.

#### Lcn2

Lcn2 is upregulated 3.54-fold by SCF+G-CSF in aged APP/PS1 mice in our study (Table 1). Also known as neutrophil gelatinase-associated lipocalin (NGAL), Lcn2 belongs to the lipocalin protein family^29^ which comprises a diverse group of transporter proteins involved in metabolism, cell growth, immune regulation, prostaglandin synthesis, iron (Fe^2+^) transport, and cell proliferation and migration.^29,30^ Lcn2 is also upregulated in various cancers^29–32^ and is a known component of neutrophil granules,^33^ where it facilitates matrix metalloproteinase 9 activity, contributing to extracellular matrix (ECM) degradation.^30^

Although some studies suggest that Lcn2 exerts primarily harmful effects, the literature remains divided.^31,34^ Lcn2 has been found to be highly expressed in the brains of AD patients and vascular dementia patients.^29,35,36^ One study reported that Lcn2 may exacerbate neuronal damage in AD, ^29^ while another study suggested that Lcn2 may exert neuroprotective effects, including memory preservation, in response to systemic inflammation.^31^ Interestingly, Lcn2 knockout (KO) mice exhibit elevated peripheral pro-inflammatory cytokines and impaired behavioral phenotypes compared to WT mice.^31^

Another study demonstrated that Lnc2 KO mice had increased susceptibility to *E. Coli* infection and reduced bacteriostatic activity.^37^ In bacterial infection, LPS stimulation of macrophage cell surface receptors triggers apoptosis and cell shrinking; in this context, Lcn2 acts as an acute-phase reactant^29,31,33,38^ promoting iron sequestration to limit bacterial access to metal ions, a process thought to be relevant to AD pathogenesis.^33,34,39^

These diverse functions implicate Lcn2 as a modulator of immune responses in AD. Lcn2 is secreted by macrophages^30^ and helps recruit innate immune cells, such as myeloid cells, into the brain.^40^ This immune cell infiltration, possibly mediated by IL-17-induced activation of S100A8/A9, may represent a crucial component of the brain’s response to amyloid pathology in aged APP/PS1 mice (Figure 4, S2).

When evaluating a gene such as *Lcn2*, both its potentially harmful and beneficial functions must be considered. Pro-inflammatory signaling is essential for responding to immune system challenges, whether caused by pathogens or Aβ accumulation. Our study supports the notion that the proinflammatory role of Lcn2, as part of an SCF+G-CSF-upregulated gene cluster, is integral to the host’s response to amyloid burden (Figure 4). Like S100A8/A9, Lcn2 mediates anti-bacterial responses as part of the neutrophil granule machinery, reinforcing the idea that inflammation is not inherently detrimental but rather necessary for immune regulation and surveillance. These findings challenge the traditional neuroinflammatory model, suggesting that a certain degree of neuroinflammation may be required to effectively counteract AD pathology in the brain.

#### Ngp

Ngp is upregulated 3.19-fold by SCF+G-CSF in aged APP/PS1 mice in our study (Table 1). Ngp protein is typically found in the cytoplasm of neutrophilic precursor granules^41^ and myeloid cells of the bone marrow.^41,42^ It is highly regulated by colony-stimulating factor 2 (CSF2 or GM-CSF), upregulated during early bone marrow development to promote myeloid differentiation, yet downregulated during granulocyte maturation^41^ to prevent excessive activation. Thus, our study’s finding that SCF+G-CSF upregulates Ngp is consistent with its known immune regulatory role.

Ngp is linked to NETosis, a process where neutrophils release extracellular traps composed of chromatin, histones, and granules to immobilize pathogens.^43^ NETosis is triggered by inflammatory cytokines and pathogens and activates NAPDH oxidase system, essential for pathogen and microbial clearance.^43,44^ Although NETosis contributes to vascular oxidative stress in AD ^45,46^, it may also reflect an immune attempt to target amyloid.

Our data suggest that SCF+G-CSF-induced Ngp upregulation may enhance amyloid recognition and clearance. Ngp is considered a microglial “sensome” gene, involved in activation and Aβ localization.^47^

#### Ltf

*Ltf* is upregulated 2.32-fold by SCF+G-CSF treatment in aged APP/PS1 mice (Table 1). Also known as lactotransferrin, it plays roles in Fe^2+^ sequestration, immune modulation, antimicrobial, anti-viral antioxidant, anti-inflammatory, and even anti-cancer functions.^48^ Ltf is expressed on small vessels near neurofibrillary tangles, amyloid deposits, and senile plaques,^49^ supporting our postulation that the UDEGs cluster (Figure 4) promotes inflammatory-mediated amyloid localization.

Ltf transport across the blood-brain barrier is thought to be enhanced in inflammatory and neurodegenerative processes, such as AD;^50^ however, it is also expressed by activated macrophages.^49^ This finding further emphasizes that activated macrophages require inflammatory signal stimulation. Notably, lactoferrin administration has been shown to reduce Aβ plaque formation and improve spatial learning in APP/PS1 mice.^51^ The functions of this gene require further explored, and its potential benefits should not be overlooked.

#### Camp

Camp is upregulated 2.01-fold by SCF+G-CSF treatment in aged APP/PS1 mice (Table 1). It is part of the 6-membered upregulated cluster, which we believe plays a role in reducing the AD state in these mice (Figure 4). Camp, not to be confused with the signaling molecule cAMP, codes for the cathelicidin antimicrobial peptide or cathelicidin-related antimicrobial peptide, including its human homologue, LL-37.^52^ These proteins are key modulators of the innate immune system.^52^

In LPS-mediated neuroinflammation, Camp is upregulated and expressed in microglia, astrocytes, and surrounding neurons, making it a potential therapeutic target for inflammatory conditions.^52^ The roles of Camp and the other genes in the 6-membered cluster, suggest that additional pathologies may contribute to AD, warranting further investigation into these diverse players in AD neuropathology.

Beyond genetic and aging factors, viral and bacterial infection, and the gut microbiome may accelerate AD progression.^53^ A study showed that Herpes Simplex Virus (HSV) in the brain increased AD risk by1.3-fold, and APOE4 mutation with HSV infection raised the risk 2.7-fold.^54^ Oral bacterial infections have also been linked to increase Aβ_42_ generation in the brain.^53^ Infections produce reactive oxygen species (ROS) and neurotoxic factors like TNFα^54^, activating the innate immune system.^55^

Antimicrobial peptides such as alpha- and beta-defensins^55^, along with Camp, can be induced by vitamin D3^55,56^ and are highly expressed in myeloid cell lines like bone-marrow derived macrophages, especially in hematopoietic diseases such as acute myeloid leukemia.^55^ Camp and other antimicrobial peptides are also involved in wound healing via MAPK, ERK, and Janus Kinase signaling pathways.^57^ Camp plays a key role in cellular regeneration, promoting cell migration, proliferation, and neovascularization^58^. Therefore, SCF+G-CSF may enhance Camp functions to reduce AD neuropathology through multiple immunomodulatory pathways, activating the immune response while promoting cell repair and regeneration.

### 3.3. Functions and Implications for SCF+G-CSF – Down-regulated Differentially Expressed Genes (DDEGs)

The top 5 known DDEGS are shown in Figure 2B and Table 1. Their functional implications in SCF+G-CSF-treated aged APP/PS1 mice are derived from transcriptomic analysis of gene chip data and a literature review.

#### Gprin3

The *Gprin3* is the most significantly downregulated DEGs in our dataset, with is a 4.60-fold decrease by SCF+G-CSF treatment in aged APP/PS1 mice (Table 1). The Grpin3 gene encodes G-protein-regulated inducer of neurite outgrowth (GPRIN) 3 protein (GPRIN3). GPRIN family proteins are highly expressed in inhibitory medium spiny neurons (MSNs) in the striatum, where they modulate dopamine type 2 receptor functions and ensure the proper sorting of MSN.^59^ These functions are implicated in goal-direct behaviors and movement disorders such as addiction, Parkinson’s disease, Huntington’s disease, and a wide range of psychiatric disorders.^59^

GPRIN3 acts as a neuromodulator, serving as an intermediary between G-protein-coupled receptors and intracellular messengers that regulate neurite growth.^59^ It is thought to exert this effect through the beta-arrestin-2 Akt pathway.^59,60^ Cas9/CRISPR knockout of the Grpin3 gene, resulted in increased dendritic arborization and decreased neuronal excitability.^59^ Notable, our previous study showed that SCF+G-CSF treatment restores dendritic loss in the cortex and hippocampus of aged APP/PS1 mice.^11^ The robust reduction in Gprin3 gene expression induced by SCF+G-CSF treatment in aged APP/PS1 mice, as shown in the current study, may be essential for SCF+G-CSF-enhanced dendritic regeneration.

#### Kcnq1ot1

The *Kcnq1ot1* gene is downregulated 2.68-fold by SCF+G-CSF in aged APP/PS1 mice (Table 1). This gene encodes a long non-coding RNA (lncRNA) rather than a protein, and it is known to promote cancer progression by regulating cell growth and proliferation through transcriptional activation and repressing protein coding genes.^61,62^ As a result, Kcnq1ot1 is involved in diverse processes such as cell cycle regulation, migration, invasion, angiogenesis, and the immune response to cancer.^61,62^

Located on the antisense strand of the Potassium Voltage-Gated Channel Subfamily Q Member 1 (KCNQ1) protein-coding gene,^62^ Kcnq1ot1 has been implicated in several solid tumors (e.g., breast, lung, and gastrointestinal cancers) as well as in hematologic cancers, particularly Acute Promyelocytic Leukemia (APML).^61^ It is also being explored as a biomarker for neurological and developmental disorders, including AD.^63–65^

Ion channels such as KCNQ1 are increasingly recognized for their roles in cancer, partly through activation of Wnt/β-catenin signaling pathway, a key oncogenic mechanism.^66^ Given that APML is characterized by uncontrolled proliferation of immature granulocytes in the bone marrow^61^, the downregulation of Kcnq1ot1by SCF+G-CSF treatment in aged APP/PS1 mice may reflect a shared anti-cancer and neuroprotective mechanism. These findings are unexpected and intriguing, suggesting that SCF+G-CSF treatment may influence overlapping pathways in cancer and neurodegeneration, warranting further investigation.

#### Akap9

The *Akap9* gene is downregulated 2.50-fold by SCF+G-CSF treatment in aged APP/PS1 mice (Table 1). Akap9 encodes A-kinase anchor protein 9, which is involved in T-cell polarization, migration, and interactions with antigen-presenting cells.^67,68^ It has been reported that AKAP9 deletion in CD4^+^ and CD8^+^ T cells does not impact T cell priming, expansion, or migration to inflamed tissue sites, but does reduce reactivation and retention at these tissues.^67^

Like Kcnq1ot, Akap9 upregulation has been associated with acute myeloid leukemia and may act through the Wnt/β-catenin pathway, a key oncogenic signaling cascade.^69^ Thus, its downregulation by SCF+G-CSF may reflect anti-cancer potential of the treatment, as supported by our analyses showing SCF+G-CSF-induced negative regulation of MET activity (Figures 4, S3).

Notably, rarer variants of Akap9 have even been linked to increased AD risk in African Americans (AAs), prompting investigation of this gene as a potential AD biomarker in this population.^68–70^ One study found that AAs with AD had lymphoblastic cell lines with Akap9 gene mutations, leading to significantly altered tau phosphorylation,^68,71^ independent of APOE genotype or amyloid precursor protein expression.^68^ These findings are particularly relevant given the underrepresentation of AAs in AD research, despite their more than twofold higher incidence compared to non-Hispanic whites.^68,70^

Together, these data suggest that Akap9 downregulation by SCF+G-CSF is a promising finding with potential implications for both AD and cancer, particularly in diverse populations.

#### Cbl

The Cbl gene is downregulated 2.42-fold by SCF+G-CSF treatment in aged APP/PS1 mice (Table 1). Cbl encodes Casitas B-Lineage Lymphoma protein, a member of the RING finger ubiquitin ligase family, which functions as both a positive and negative regulator of signal transduction via receptor- and non-receptor kinases.^72–74^ It also negatively regulates signaling pathways in hemopoietic and immune cells.^73^

Like Kcnq1ot1, Cbl is associated with hematologic malignancies, and its overexpression induces myeloid leukemia in mice.^72,74^ The downregulation of cancer-promoting pathways involved in Cbl by SCF+G-CSF treatment may help shift the immune landscape toward HSC-mediated immune function. This is supported by TAC pathway analysis, which showed Cbl downregulation in multiple signaling pathways, including IL-7, IL-2, IL-4, and EGFR1 (Figure 5). Our study found Cbl to be downregulated in over 40% of enriched pathways identified by Reactome pathway analysis (Figure 6), with its regulatory functions showing approximately 50-fold enrichment following SCF+G-CSF treatment.

**Figure 5.**
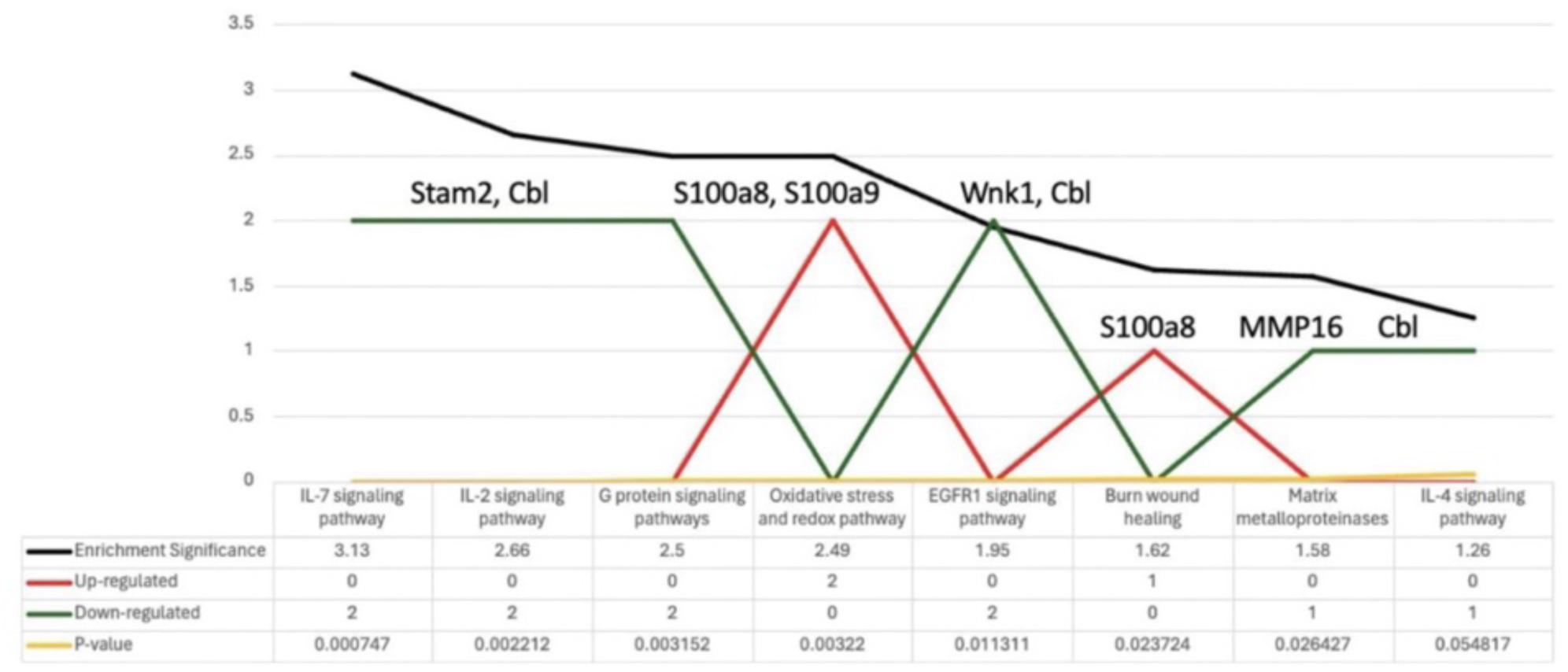
Transcriptome Analysis Console (TAC) pathway enrichment analysis of SCF+G-CSF-induced differentially expressed genes (DEGs) in the brains of aged APP/PS1 mice. SCF+G-CSF treatment upregulates pathways related to S100a8/a9-mediated oxidative stress and wound healing. SCF+G-CSF-downregulated DEGs reduce inflammatory pathways, EGFR1 signaling, and MMP16 activity.

These findings emphasize the complex overlap between cancer and neurodegeneration, particularly through shared mechanisms involving immune modulation. Cancer cells depend on hijack cell machinery to proliferate, evade immune surveillance, and invade healthy tissues, often by suppressing the tumor microenvironment.^75^ In contrast, SCF+G-CSF treatment appears to stimulate immune function, potentially counteracting cancer-associated immunosuppression.

**Figure 6.**
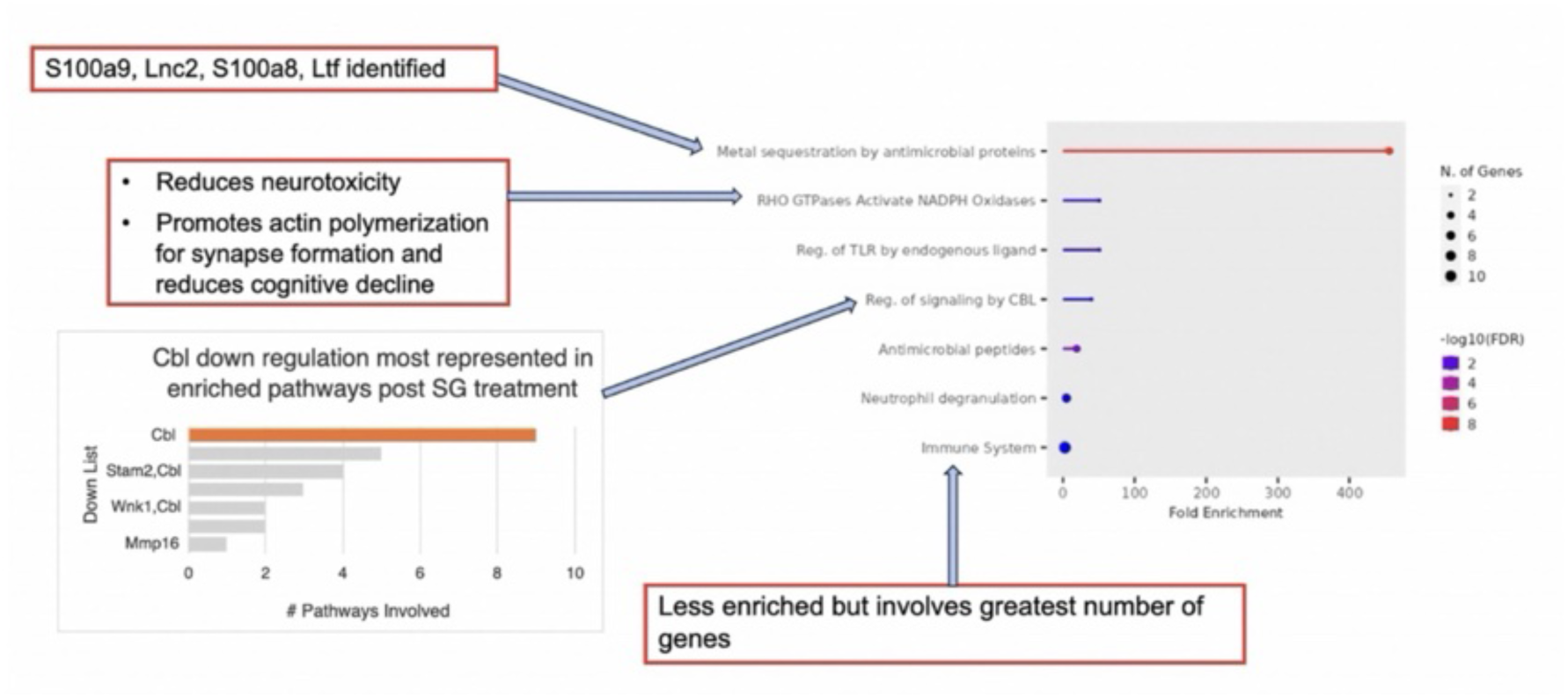
Reactome pathway analysis of SCF+G-CSF-induced differentially expressed genes (DEGs) in the brains of aged APP/PS1 mice. SCF+G-CSF treatment exerts diverse effects in aged APP/PS1 mice, including reduction of AD pathology and suppression of key cancer­promoting genes. The most enriched pathway involves metal ion sequestration, suggesting potential chelation activity of SCF+G-CSF. Additional enriched pathways are associated with reduced neurotoxicity, enhanced synapse formation, improved cognitive function, immune system activation, and downregulation of *Cbi,* a major cancer-promoting gene, which appears in over 40% of the enriched pathways.

Cancer and neurodegeneration are linked by their multifactorial nature from an epidemiological standpoint, as well as by overlapping mechanisms involving inflammation, DNA damage, oxidative stress, and various forms of metabolic dysfunction.^75,76^ A large cohort study of over 1,200 participants found that those diagnosed with AD were less likely to develop cancer, and cancer survivors had a reduced risk of developing AD compared to those without cancer.^75^ SCF+G-CSF’s anti-cancer potential is supported by its ability to downregulate Cbl, as well as other cancer-promoting genes like Kcnq1ot1 and Akap9. These findings suggest that reducing shared mechanisms of cancer and neurodegeneration, including inflammation, DNA damage, oxidative stress, and metabolic dysfunction, may be critical to improving outcomes in AD.

#### Snhg11

The Snhg11 gene is downregulated 2.41-fold by SCF+G-CSF treatment in aged APP/PS1 mice (Table 1). Also known as *Sonic Hedgehog 11*, this long non-coding RNA (IncRNA) is differentially expressed in our dataset. Along with *Kcnq1ot1* and *Akap9*, *Snhg11* is known to activate the cancer-promoting Wnt/β-catenin signaling pathway,^77^ and its upregulation has been associated with poor cancer prognoses.^69,77,78^ *Snhg11* has been implicated in oncogenesis through the activation of Notch3.^79^ Notably, mutations in Notch3 have been identified in Cerebral Autosomal Dominant Arteriopathy with Subcortical Infarcts and Leukoencephalopathy (CADASIL), the most common inherited form of vascular dementia.^79^ Additionally, *Snhg11* has been linked to cognitive decline in Down syndrome through its stabilization of hypoxia-inducible factor^80^ highlighting the broader relevance of lncRNA in both intellectual disability and tumor metastasis.^81^

### 3.4. SCF+G-CSF treatment induces interaction and co-expression of 6-membered gene cluster to reduce AD neuropathology in APP/PS1 mice

Our data demonstrate that the transcriptomic profile induced by SCF+G-CSF is highly specific and fine-tuned, as reflected in the functional significance of differentially expressed genes (DEGs) that directly address AD neuropathology. STRING v12.0 interaction analysis reveals that SCF+G-CSF treatment enhances both the interaction and co-expression of a 6-member gene cluster including S100a8, S100a9, Lcn2, Ngp, Ltf, and Camp (Figure 4). While 19 of the 27 DEGs in this dataset are downregulated by SCF+G-CSF, this specific gene cluster includes 8 of the remaining upregulated DEGs induced by SCF+G-CSF, with two genes lacking current annotation (Figure 2). This observation suggests that SCF+G-CSF treatment in aged APP/PS1 mice exerts a targeted effect on AD-related pathways rather than broadly altering gene expression, supporting its potential tolerability in AD patients. Notable, Sargramostim, the human recombinant form of GM-CSF,^5^ a hematopoietic growth factor, has already been reported to be safe and effective in AD patients.^5^

The interaction and co-expression within this gene cluster have been shown to enhance microglial function, contributing to improved AD-related outcomes in APP/PS1 mice. Specifically, S100A8 and S100A9 are known to facilitate amyloid localization, tagging, and subsequent clearance via phagocytosis and degradation by innate immune cells.^20,21,23^ These two genes were identified as the most upregulated DEGs in response to SCF+G-CSF treatment, and their co-expression correlation was remarkably high at 0.999, indicating near-complete co-expression (Figure 4).

Understanding the functions and spatial expression of the genes in this cluster is important. STRING interaction analysis reveals that this 6-member gene cluster is functionally linked to the spleen and immune system (Figure 4). The spleen, a secondary lymphoid organ, like lymph nodes, plays a key role in filtering blood and orchestrating immune responses. A previous study reported that 40-60% of amyloid burden is found in the peripheral system, and that splenectomy exacerbates AD pathogenesis in APP/PS1 mice by increasing amyloid beta accumulation and worsening cognitive deficits.^82^ These findings reinforce the critical role of peripheral immune dysregulation in AD progression. In the present study, the SCF+G-CSF-upregulated 6-member gene cluster was detected in brain tissue from aged APP/PS1 mice, suggesting that SCF+G-CSF may mitigate AD neuropathology by enhancing immune system function and promoting coordination between central and peripheral immune responses.

### 3.5. IL-17 may drive the expression of S100a8, S100a9, and Lcn2 genes to promote innate and adaptive immune cell infiltration into the brains of aged APP/PS1 mice

ShinyGO 0.77 Analysis reveals that SCF+G-CSF induces 29.1-fold enrichment of the IL-17 signaling pathway, identifying 3 genes (*S100a8, S100a9, and Lcn2)* from our dataset as part of this pathway (Figure S2). These findings are corroborated by STRING v12.0 interaction analysis (Figure 4), which also supports the functional interaction of these genes. Based on our data and prior literature,^83^ we postulate that the SCF+G-CSF treatment-induced upregulation (Figure 2) and interaction of S100a8, S100a9, and Lcn2 is stimulated by IL-17, facilitating innate and adaptive immune cell infiltration into the brains of aged APP/PS1 mice (Figure S2).

This immune activation may be necessary for mitigating AD-related neuropathology. However, the role of IL-17 in AD remains controversial. While some findings suggest that IL-17 promotes neuroinflammation, this inflammatory response is also necessary for glial production of chemokines that attract myeloid cells and T cells.^83^ It is thus unsurprising that anti-IL-17 (and anti-IL-23) antibodies have been proposed as potential AD treatments^83^, and their ablation was shown to reduce CD8+ T cell and myeloid cell infiltration into the brain in APP/PS1mice.^83,84^ Despite these efforts, eliminating CD8⁺ T cells from the brain, spleen, and blood of APP/PS1 mice has not led to improvements in cognitive function or changes in amyloid plaque burden.^84^

This raises an important question: If neuroinflammation is a hallmark of AD, and IL-17 promotes this process, why does not IL-17 ablation attenuate AD pathology? Our study offers a potential explanation. While IL-17 promotes immune cell infiltration, this may support beneficial immune responses such as amyloid localization and clearance. Our STRING analysis (Figure 4) implicates S100a8, S100a9, and Lcn2 as participants in IL-17 signaling. We propose that IL-17 may stimulate these genes to promote immune cell infiltration into the CNS, aiding in the clearance of amyloid deposits.

Taken together, supported by both ShinyGO (Figure S2) and STRING (Figure S3) analyses as well as our previous studies,^12,85^ our findings suggest that SCF+G-CSF treatment enhances immune cell infiltration into the brain of aged APP/PS1 mice, potentially reducing AD neuropathology through an IL-17–mediated mechanism. An alternative interpretation, however, is that IL-17 signaling may not be directly induced by SCF+G-CSF but rather represents a key component of AD pathology that SCF+G-CSF is insufficient to suppress. Future studies should explore IL-17 pathway subtypes and their regulation to clarify both the mechanism of SCF+G-CSF action and the broader role of immune modulation in AD pathogenesis.

### 3.6. Transcriptomic analysis reveals that SCF+G-CSF downregulates EGFR1, interleukin signaling pathways, and matrix metalloproteinase 16 to reduce amyloid and NFT formation

STRING v12.0 interaction analysis identified *Cbl* and *Stam2* (signal transducing adaptor molecule) as interacting genes downregulated by SCF+G-CSF treatment (Figures 4 and 5, Table 2). Both genes are involved in the epidermal growth factor (EGF) signaling pathway. *Stam2*, which is ubiquitously expressed across tissues, binds to Janus kinases (JAKs), whose activity is dependent on cytokines such as IL-2 and GM-CSF. These interactions regulate DNA synthesis and induce expression of the c-Myc oncogene.^86^ JAKs bind to the common γc and βc receptor chains, which are shared by interleukins 2, 3, 4, 5, 7, 9, 15, and GM-CSF.^86^ Upon stimulation by ligands, including platelet-derived growth factor (PDGF) and EGF, these receptors activate downstream targets such as *Stam2* via tyrosine phosphorylation.^86^

**Table 2.**
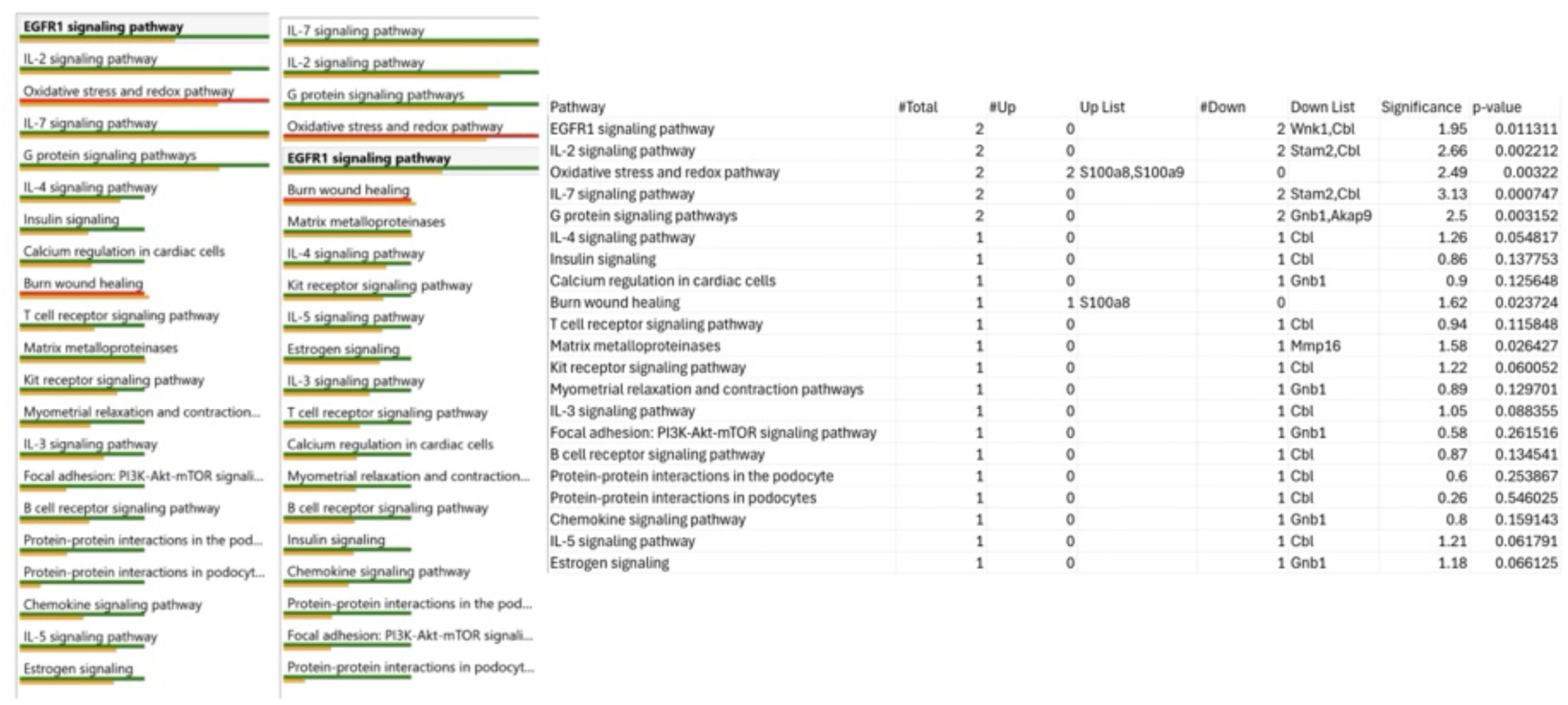
SCF+G-CSF-induced pathway enrichment in the brains of aged APP/PS1 mice.

In our study, SCF+G-CSF led to the downregulation of IL-2, IL-4, IL-7, and epidermal growth factor receptor 1 (EGFR1) (Figure 5 and Table 2). EGFR1 is a well-characterized member of the receptor tyrosine kinase family, which plays a central role in cell proliferation, survival, and migration.^87,88^ EGFR1 is commonly overexpressed in cancer,^87^ and emerging evidence suggests its overactivation is also relevant to AD pathogenesis. Specifically, preformed amyloid fibrils can stimulate EGFR1, promoting further amyloid aggregation^89,90^ and tau phosphorylation.^90^

Collectively, our findings and previous studies suggest that SCF+G-CSF treatment attenuates AD pathology by downregulating EGFR and interleukin signaling pathways, thus reducing the neurotoxic impact of cytokine dysregulation characteristic of the disease.

Additionally, SCF+G-CSF treatment was found to downregulate the matrix metalloproteinase 16 (MMP16) pathway, which may further contribute to reduced AD neuropathology (Figure 5, Table 2). MMPs are zinc- and calcium-dependent enzymes that degrade components of the extracellular matrix (ECM), including collagen, elastin, gelatin, and glycoproteins^91^. The ECM provides structural and biochemical support to neurons and serves as a protective barrier around neurons. In pathological states, MMPs contribute to inflammation and neurodegeneration by remodeling the perineuronal environment, altering the distribution of neurotransmitter and growth factor receptors.^91^ Members of the MMP family have been detected in senile plaques, neurofibrillary tangles, and in hippocampal and cortical neurons, and their expression is elevated in AD patients, where they are implicated in cognitive decline.^91^ Based on our data and corroborating evidence, we propose that SCF+G-CSF downregulates MMP16 to limit ECM degradation, thereby reducing neuronal vulnerability and potentially improving cognitive outcomes in AD.

### 3.7. Metal ion chelation and immune modulation may contribute to SCF+G-CSF-reduced AD neuropathology

The role of metal ion dys-homeostasis in AD neuropathology is increasingly being explored. Emerging evidence suggests that metal ions such as zinc, manganese, copper, and iron not only enhance amyloid-β production but also promote amyloid-tau aggregation.^92–94^ It is important to consider that metal-induced cell death pathways, including ferroptosis (iron-mediated) and cuproptosis (copper-mediated), may contribute to AD neuropathology such as neuroinflammation and cognitive decline.^93^

Evidence linking impaired metal ion regulation to the neuroinflammatory pathology of AD has led to the proposition that metal chelators could serve as potential therapeutic agents.^93^ In addition to their role in inflammation and protein aggregation, metal ions contribute to oxidative stress, neuronal loss, tau hyperphosphorylation, neurofibrillary tangle formation, and free radical generation—particularly through copper–amyloid complexes, which can lead to severe cellular damage.^94^

Although the metal ion hypothesis of AD remains under debate, our findings provide novel support for its relevance to both AD pathology and the mechanism of SCF+G-CSF treatment in APP/PS1 mice. Our analysis revealed over 400-fold enrichment of the metal ion sequestration by antimicrobial proteins pathway (Figure 6), and a STRING interaction strength of 3.08 among four co-expressed genes (S100a8, S100a9, Ltf, and Lcn2) that mediate this function (Figure 4). These findings strongly suggest that SCF+G-CSF may act as a metal chelator, promoting metal sequestration to restrict ion availability and thereby preventing their involvement in harmful AD-related processes.

In addition to metal chelation, our study suggests that intracellular and extracellular transport dysfunction should be considered an integral component of AD neuropathology. The role of Rho GTPases, which belong to the RAS superfamily and regulate cargo transport within and outside cells, ^95^ is gaining increasing attention in AD research. Rho GTPase dysfunction has been associated with Aβ plaque and NFT formation, as well as neurotoxicity.^95^ Interestingly, SCF+G-CSF treatment specifically enriched Rho GTPase–associated pathways that activate NADPH oxidases (Figures 4, 6, and S3), a function known to reduce free radical accumulation and neurotoxicity.

This SCF+G-CSF-induced enrichment of Rho GTPase signaling (Figure S3) may represent a mechanism through which the treatment improves cognitive function, as Rho GTPase activity has been shown to promote actin polymerization, synapse formation, and behavioral improvements in AD mouse models.^95^

Two major activators of Rho GTPases, Ras and EGFR are implicated in AD pathology. Notably, EGFR1 was downregulated by SCF+G-CSF treatment (Figure 5), suggesting that in addition to its anti-cancer effects, SCF+G-CSF may suppress EGFR1 to reduce Rho GTPase–driven tau pathology.^95^

However, the concurrent enrichment of Rho GTPase functions specific to NADPH oxidase activation highlights a selective mechanism by which SCF+G-CSF supports oxidative burst activity, a process crucial for degradation of intracellular and extracellular tau and amyloid.^20,23^

### 3.8. A significantly greater number of genes are upregulated in APP/PS1 mice in the absence of SCF+G-CSF treatment

Given that our previous studies demonstrated that SCF+G-CSF treatment reduced AD neuropathology in aged APP/PS1 mice^11,12^, we sought to use transcriptomic analysis to determine how advanced-stage AD alters gene expression in aged APP/PS1 mice. To accomplish this, we compared gene chip (microarray) data from three 17–18-month-old APP/PS1 mice to that of two age-matched WT control mice. Both groups received subcutaneous vehicle injections for 12 consecutive days and were euthanized on day 13. RNA was extracted from the frontal lobes for transcriptomic analysis (Figure 1). Again, we specifically selected aged APP/PS1 mice to represent a more progressed or chronic stage of AD, which we believe better reflects the human AD patient population.

A list of DEGs in APP/PS1 (CON) mice compared to WT mice is provided in Table 3 and Figure 7. A heat map illustrating hierarchical clustering of the DEGs is presented in Figure 8, where red indicates upregulated genes and blue indicates downregulated genes.

**Table 3.**
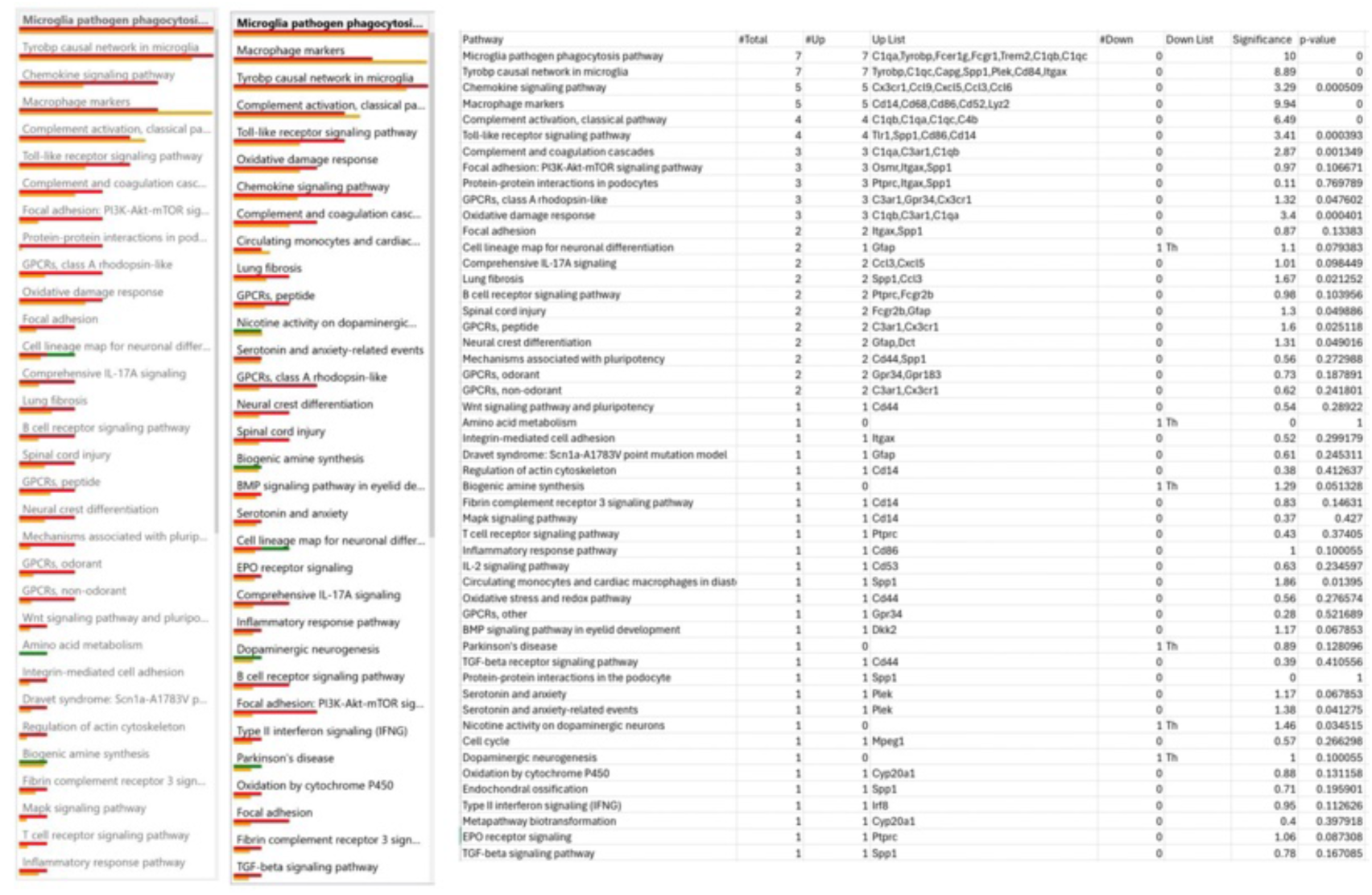
Pathway Enrichment in APP/PS1 (CON) vs WT mice in the absence of SCF+G-CSF.

**Figure 7.**
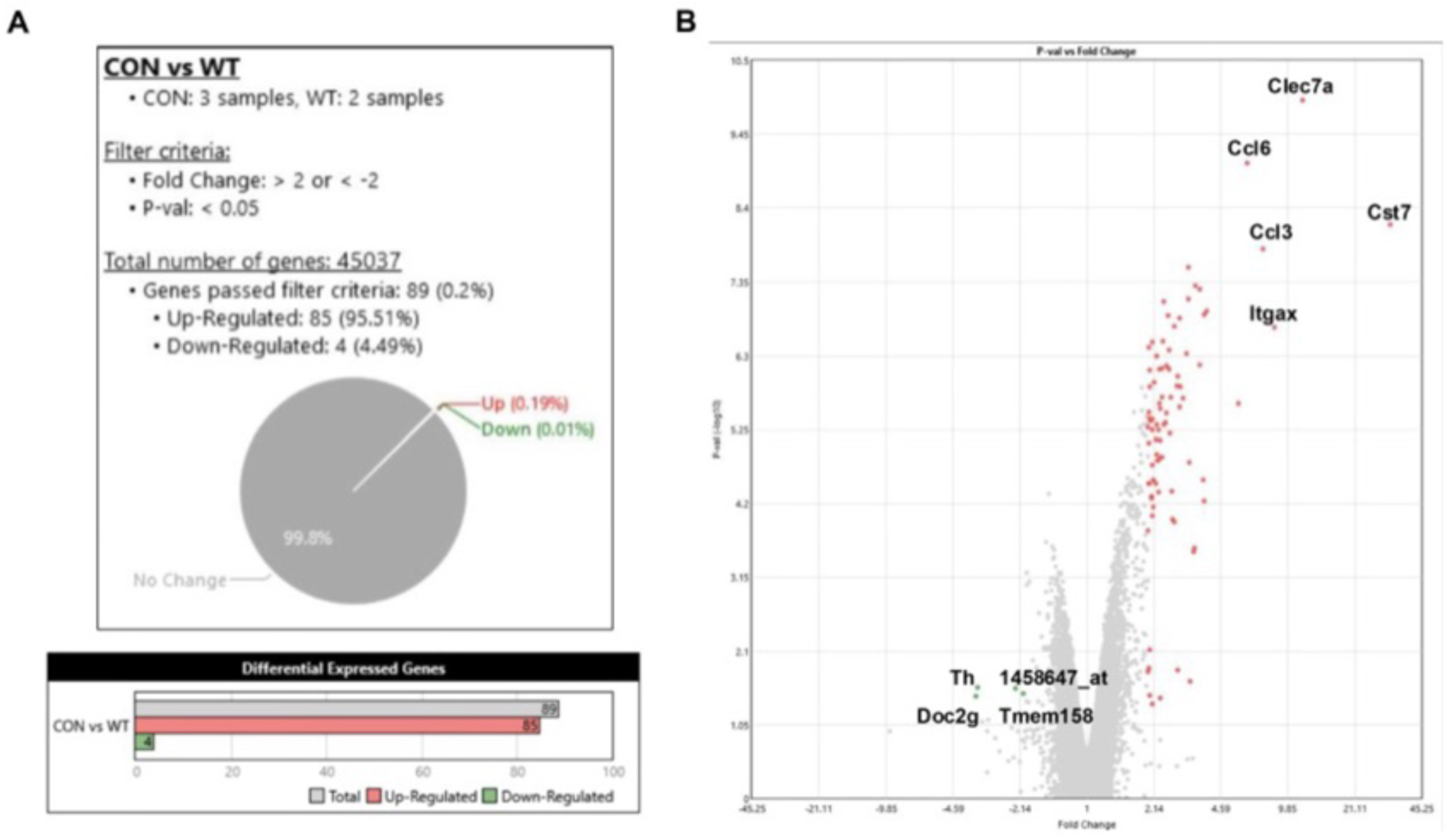
Summary and volcano plot of gene expression differences between aged APP/PS1 and aged WT mice analyzed using the Transcriptome Analysis Console (TAC). (A) A total of 89 differentially expressed genes (DEGs) were identified in the brains of aged APP/PS1 mice compared to age-matched WT controls, with over 95% of the DEGs upregulated. This indicates a predominance of gene upregulation in APP/PS1 mice lacking SCF+G-CSF treatment. (B) The top five most significantly upregulated and dowπregulated DEGs are labeled and bolded. Genes labeled only by numerical probe set identifiers lack known gene names or symbols. CON: aged APP/PS1 mice injected with vehicle; WT: age-matched wild-type mice.

**Figure 8.**
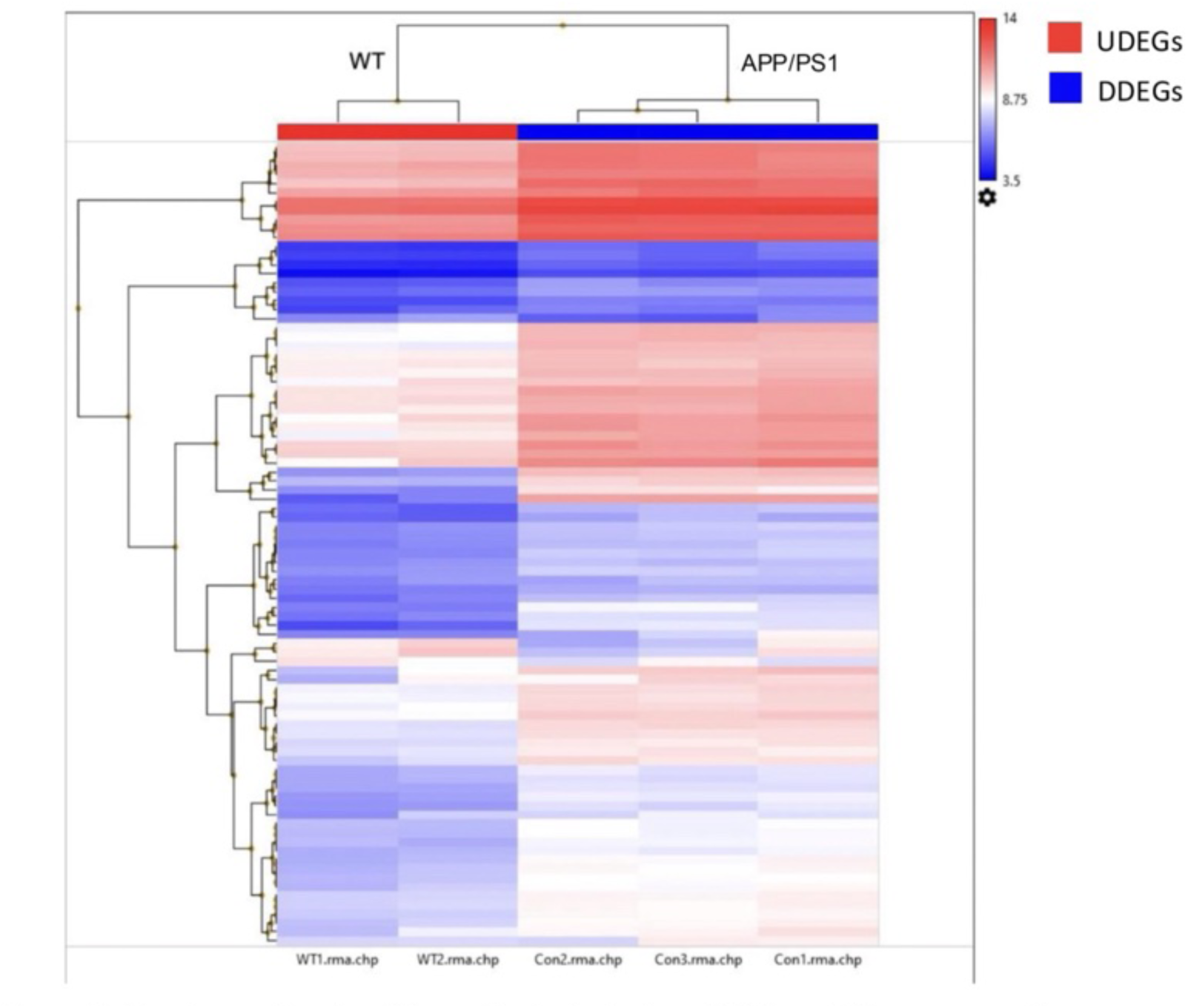
Heat map showing hierarchical clustering of differentially expressed genes (DEGs) in aged WT and APP/PS1 mice. Normalized gene expression is displayed on a log-scaled gradient, where darker colors indicate higher expression levels and lighter colors indicate lower expression levels. Upregulated DEGs are marked in red, and downregulated DEGs in blue. The heat map was generated using the Transcriptome Analysis Console (TAC). CON: aged APP/PS1 mice injected with vehicle; WT: age-matched wild-type mice.

Gene expression analysis using ThermoFisher Scientific’s Transcriptional Analysis Console revealed that, out of the 45,037 genes detected, 89 genes (0.2%) were significantly differentially expressed (≥2-fold change) in APP/PS1 mice compared to WT controls (Figure 7). Among these 89 DEGs, 85 genes (95.5%) were upregulated (UDEGs), and 4 genes (4.5%) were downregulated (DDEGs) (Figure 7).

In contrast to this finding, SCF+G-CSF treatment led to downregulation of more than 70% of DEGs, with approximately 29.6% of DEGs being upregulated in aged APP/PS1 mice (Figure 2). These findings suggest that, in the absence of SCF+G-CSF treatment, gene expression in aged APP/PS1 mice is less transcriptionally controlled compared to WT mice (Figure S4).

### 3.9. Transcriptional analysis reveals uncontrolled, upregulated amyloid clearance mechanisms and microglial activation in APP/PS1 mice compared to WT controls

Next, we sought to identify the biological pathways associated with our DEG dataset. Interestingly, we found that cell functions relevant to AD treatment were already upregulated in untreated APP/PS1 mice compared to healthy WT controls (Figure 9, 10A, 10B).

**Figure 9.**
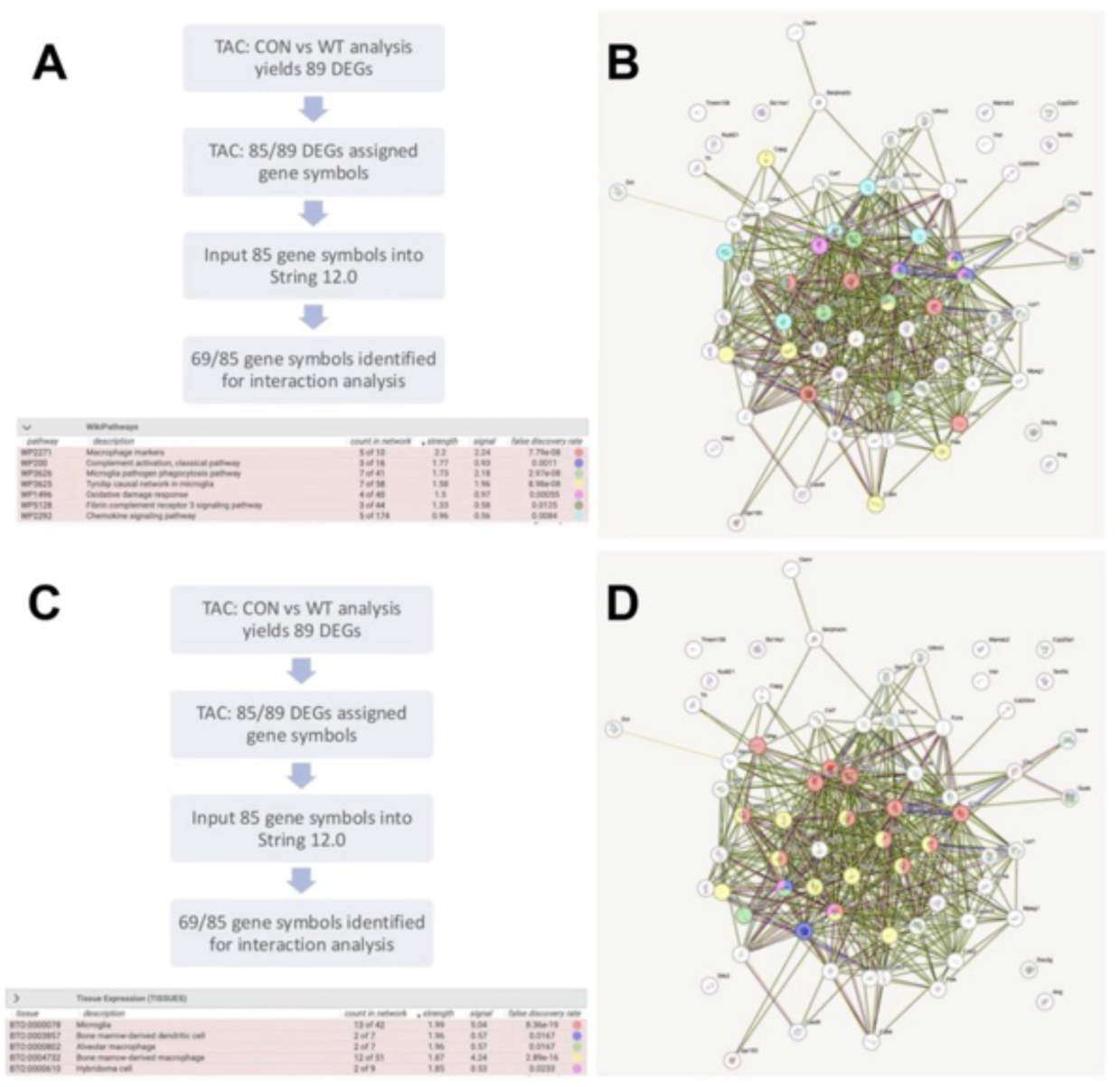
STRING v12.0 pathway analysis of differentially expressed genes (DEGs) highlights enrichment in macrophage markers and innate immune system pathways in aged APP/PS1 mice. (**A**, **C**) Transcriptome Analysis Console (TAC) identified 89 DEGs, of which 85 had known gene names. Of these 85 genes, 69 were identified by STRING v12.0 interaction analysis. Enriched functional clusters are color-coded by pathway type. (**B**) A prominent gene cluster shows the highest enrichment in macrophage marker-associated pathways and innate immune responses, including complement and phagocytosis pathways, indicating activation of immune signaling in aged APP/PS1 mouse brain in response to AD pathology. (**D**) Another enriched gene cluster includes pathways expressed by microglia and bone marrow-derived macrophages, emphasizing the contribution of both central and peripheral immune cells in the AD state. CON: aged APP/PS1 mice injected with vehicle; WT: age-matched wild-type mice.

Both String 12.0 and ATC analyses revealed that large gene cluster with the highest enrichment in macrophages, bone marrow-derived macrophages/dendritic cells, and microglia as well as these cell-involved pathways and innate immune system pathways, including microglia pathogen phagocytotic pathways, complement activation (classical) pathways, and chemokine signaling pathway (Figures 9 and 10). These data suggest that the immune system in APP/PS1 mice is activated to combat AD neuropathology. This implies that anti-AD signaling pathways and some neuroinflammation are naturally responding to AD neuropathology without intervention.

**Figure 10.**
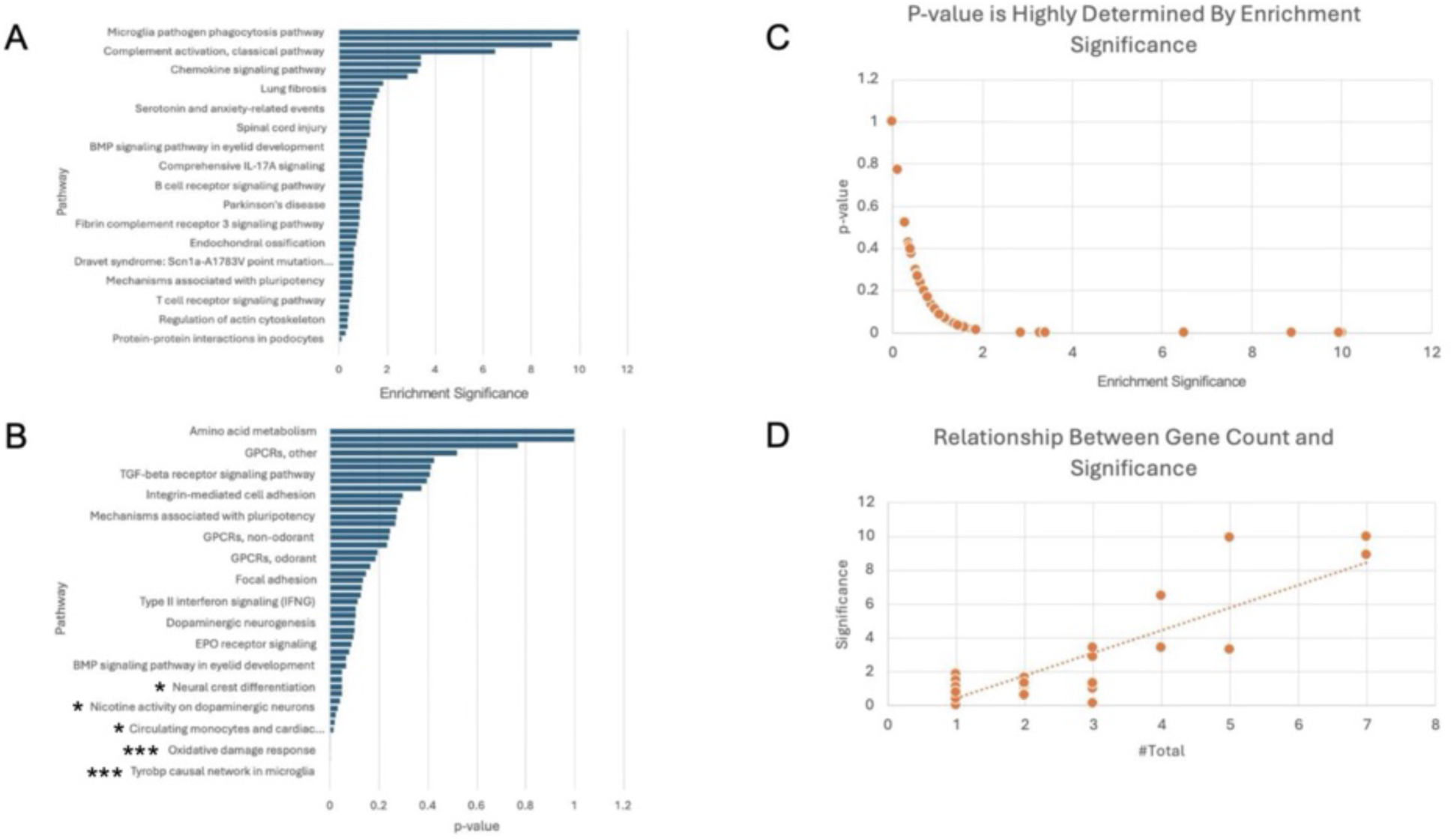
Transcriptome Analysis Console (TAC) shows enrichment of microglial and immune-related pathways in aged APP/PS1 mouse brains. (A) Pathway enrichment analysis ranked by significance highlights immune-related responses to amyloid pathology. (B) Pathway enrichment ranked by p-value; *p<0.05, **p<0.01, ***p<0.001. (C) Statistical analysis demonstrates an inverse exponential relationship between p-value and enrichment significance. (D) A positive correlation is observed between the number of differentially expressed genes (DEGs) and enrichment significance, indicating that highly enriched pathways include a greater number of DEGs.

Enriched functions in aged APP/PS1 mice, when compared to WT mice, included microglial phagocytosis, complement activation (especially the classical pathway), and chemokine signaling (Figure 10A, Table 3). Among these, the most statistically significant pathway, ranked by p-value, was the TYROBP casual microglial network, which included 7 upregulated DEGs (UDEGs): *Tyrobp, C1qc, Capg, Spp1, Plek, Cd84*, and *Itgax* (Figure 10B, Table 3). Other significantly enriched pathways involved responses to oxidative damage, as well as signatures related to circulating monocytes and cardiac macrophages associated with diastolic dysfunction (Figure 10B, Table 3).

TYROBP (Tyrosine Kinase Binding Protein) is a transmembrane immune signaling adaptor that is highly upregulated in AD patients.^96^ It plays a central role in signal transduction pathways mediated by microglia, macrophages, dendritic cells, and even osteoclasts, which can release bone-derived cells into circulation.^96^ Evidence suggests that TROBP promotes microglial phagocytosis of amyloid plaques and neuronal debris, while also exerting anti-inflammatory effects by suppressing microglia-mediated cytokine production.^96^ A previous study using PSEN1 mutant mice showed that *Tyrobp* was preferentially expressed in hippocampal microglia, and that *Tyrobp*-deficient mice exhibited slower cognitive decline.^97^ These mice also maintained learning behavior, showed reduced levels of proinflammatory cytokines (e.g., TNF-α, IL-1β, IL-6), but had fewer astrocytes and microglia in the hippocampus.^97^

Taken together, our current findings and supporting previous studies suggest that TYROBP upregulation in aged APP/PS1 mice may be a compensatory response to high amyloid burden. Interestingly, *Tyrobp* was not significantly differentially expressed in aged APP/PS1 mice treated with SCF+G-CSF compared to untreated APP/PS1 controls. This may indicate that excessive compensatory microglial activation, such as that mediated by TYROBP in AD, could be maladaptive. We propose that SCF+G-CSF treatment promotes a more regulated and beneficial level of microglial activation. The lack of TYROBP overexpression in SCF+G-CSF-treated aged APP/PS1 mice supports the idea that AD pathology is substantially mitigated by treatment, reducing the need for such compensatory mechanisms.

### 3.10. Functions and implications of the Top 5 upregulated DEGs in aged APP/PS1 mice

#### Cst7

Cst7 was markedly upregulated by 31.73-fold in aged APP/PS1 mice without SCF+G-CSF treatment (Figure 7B, Table 4). Also known as Cystatin F, Cst7 is a cysteine protease inhibitor localized to the endo-lysosome^98^ It is highly expressed in microglia and monocyte-derived macrophages during the progression of both cancer and AD.^98–100^ Additionally, Cst7 has been implicated in demyelination and remyelination, especially in the context of multiple sclerosis.^101^ Elevated levels of dimerized Cst7 have been found in the blood of AD patients,^100^ and the protein is known to bind amyloid-β (Aβ), interfering with its recognition and uptake by monocytes.^100^ This impaired clearance may contribute to amyloid accumulation and the associated inflammatory response. We postulate that the marked upregulation of *Cst7* may contribute to the activation of phagocytic pathways, such as those regulated by TYROBP, as a compensatory response in the untreated AD state. Given its roles in impaired Aβ clearance and immune dysregulation, it is not surprising that Cst7 upregulation is associated with worsening clinical signs in AD.

**Table 4.**
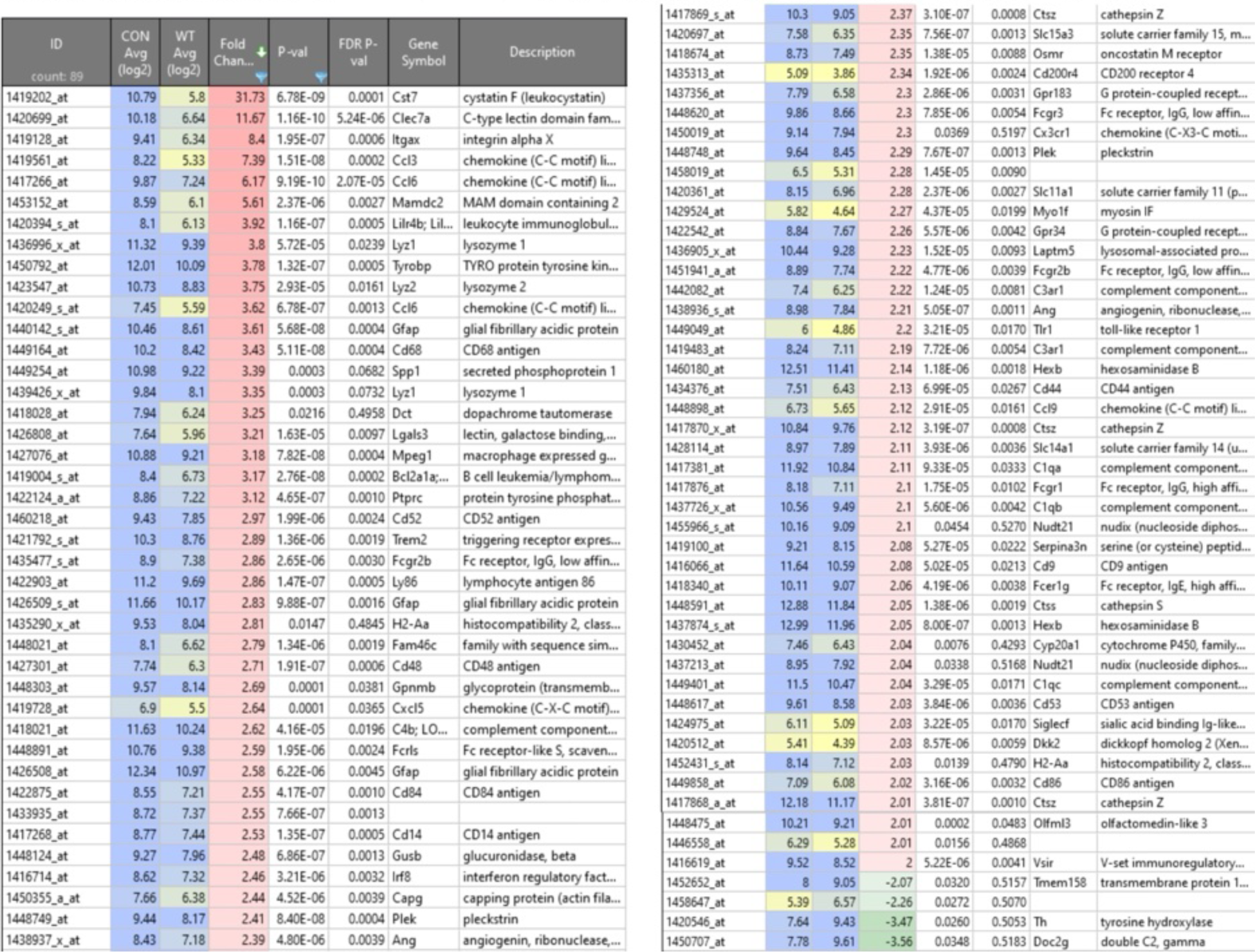
DEGs in APP/PS1 (CON) vs WT mice in the absence of SCF+G-CSF.

#### Clec7a

Clec7a was upregulated 11.67-fold in aged APP/PS1 mice without SCF+G-CSF treatment (Figure 7B, Table 4). In contrast, significantly lower Clec7a protein levels were reported in 5xFAD mice.^102^ *Clec7a* encodes a receptor expressed in neutrophils, macrophages, and dendritic cells, that recognizes beta-glucans in fungal cell walls and activates macrophage responses.^103^ It is also responsible for recruiting and activating spleen tyrosine kinase (SYK).^104^ TREM2, which our previous study showed to be upregulated by SCF+G-CSF treatment in the brains of aged APP/PS1 mice,^11,12^ is a microglial receptor that interacts with the transmembrane adaptor DAP12 to recruit SYK.^104,102,102^

These findings further support the concept that AD is primarily a disorder of immune system dysregulation. In glioma, Clec7a knockout reduced M2 macrophage polarization and chemotaxis,^105^ suggesting its role in promoting restorative immune responses.

Thus, while Clec7a inhibition may be beneficial in cancer, it could hinder protective microglial and macrophage activity in AD. Our data suggest that aged APP/PS1 mice exhibit enhanced compensatory activation of microglia and macrophages (Figures 9, 10A and 10B), highlighting the complexity of targeting Clec7a in AD therapeutics.

#### Itgax

Itgax was upregulated 8.40-fold in aged APP/PS1 mice without SCF+G-CSF treatment (Figure 7B, Table 4). Also known as Integrin subunit alpha X, or CD11c, elevated *Itgax* expression has been associated with cognitive decline and more severe AD diagnoses.^106^ In the present study, *Itgax* gene expression was significantly higher in aged APP/PS1 mice compared to WT controls. Previous work has shown that ITGAX binds to Spp1 (secreted phosphoprotein 1), which encodes osteopontin (OPN), and is involved in TYROBP signaling. CD11c⁺ microglia that produce OPN represent a distinct subtype of disease-associated microglia implicated in driving AD neuropathology.^106^

Given the advanced neuropathological stage of aged APP/PS1 mice, the upregulation of *Itgax* is expected. Importantly, its absence of upregulation in SCF+G-CSF-treated aged APP/PS1 mice supports the therapeutic potential of SCF+G-CSF, suggesting that the treatment may suppress this pathogenic microglial activation pathway.

#### Ccl3 & Ccl6

Ccl3 and Ccl6 were upregulated 7.39-fold and 6.17-fold, respectively, in aged APP/PS1 mice compared to WT controls (Figure 7B, Table 4). *Ccl3* and *Ccl6* encode the chemokine ligands CCL3 and CCL6, which act as chemo attractants for microglia and contribute to immune cell recruitment during neuroinflammatory responses.^107,108^ CCL3 secretion has been shown to be modulated by APOE in astrocytes^107^ while CCL6 may reduce detrimental neuron–astrocyte interactions, thereby protecting neurons from glutamate excitotoxicity.^108^ Additionally, CCL6 has been implicated in promoting M2 macrophage polarization at inflammatory sites via activation of the PI3K–Akt signaling pathway.^109^

### 3.11. Functions and implications of the Downregulated DEGs in aged APP/PS1 mice

#### Doc2g

Doc2g was downregulated 3.56-fold in aged APP/PS1 mice compared to WT controls (Figure 7B, Table 4). *Doc2g* encodes the double C2 domain-containing protein gamma (DOC2G), a protein belonging to the Double C2 domain family, which is typically involved in calcium-dependent binding to phospholipid membranes and regulation of synaptic vesicle exocytosis.^110,111^ While other family members, such as DOC2A and DOC2B, facilitate neurotransmitter release through phospholipid interactions, DOC2G is unique in that it does not bind phospholipids.^110^

Neurotransmitter dysregulation is a hallmark of AD, particularly involving glutamate and acetylcholine, which play essential roles in synaptic transmission, learning, and memory.^112^ The observed downregulation of *Doc2g* may reflect a compensatory response of the AD brain aimed at limiting aberrant synaptic activity or excitotoxicity during advanced stages of pathology in aged APP/PS1 mice.

#### Th

Th was downregulated 3.47-fold in aged APP/PS1 mice compared to WT controls (Figure 7B, Table 3). *Th* encodes tyrosine hydroxylase (TH), the rate-limiting enzyme in the synthesis of dopamine (DA) through the conversion of tyrosine to L-DOPA. Deficiency in *Th* is classically associated with Parkinson’s disease, where degeneration of dopaminergic neurons in the substantia nigra and loss of neuromelanin lead to motor dysfunction.^113^ This process is driven by oxidative stress, impaired autophagy, and neuroinflammation.^113^

Although the role of *Th* in AD remains under investigation, reduced TH expression has also been observed in AD brains.^114^ Amyloid-mediated damage to noradrenergic neurons, particularly those projecting to the hippocampus, has been implicated in synaptic dysfunction, neuronal loss, and cognitive decline.^114^ Thus, the observed downregulation of *Th* in aged APP/PS1 mice may result from high amyloid burden and neuroinflammatory stress, further supporting the link between AD and neurotransmission dysfunction. These findings underscore the need for continued investigation into monoaminergic deficits in AD pathogenesis.

#### Tmem158

Tmem158 was downregulated 2.07-fold in aged APP/PS1 mice compared to WT controls (Table 7B, Table 4). *Tmem158*, which encodes transmembrane protein 158, has been implicated in promoting cancer cell proliferation, migration, and survival through activation of STAT3,^115–117^ a key modulator of JAK family and EGFR signaling pathways.^115^ It may also enhance TGF-β signaling.^116,117^ While the role of *Tmem158* in AD remains poorly understood, its downregulation in our study may reflect reduced cell survival signaling and support the idea that neurodegenerative pathways are more active in the AD brain.

### 3.12. Additional functional changes implicate extracellular matrix integrity, immune dysregulation, and microbial pathologies in the aged APP/PS1 mouse brain

ShinyGO 0.77 analysis revealed that glycosaminoglycan (GAG) degradation was the most significantly enriched biological function process, showing nearly 20-fold enrichment, in aged APP/PS1 mice compared to WT controls (Figure S5). GAGs are critical components of proteoglycans,^118,119^ particularly those containing heparan sulfate (HS) polysaccharide, which maintain extracellular matrix (ECM) integrity, tissue renewal, and homeostasis.^91,119^ Notably, excessive HS GAG accumulation has been shown to promote the formation of Maltese/star-like Aβ_1–40_ deposits and tau aggregation into paired helical filaments, both of which are resistant to degradation and ultimately accumulate in the cortex and hippocampus.^118,119^

Interestingly, GAG degradation, rather than formation, was enriched in aged APP/PS1 mice without SCF+G-CSF treatment, suggesting a compensatory mechanism aimed at reducing pathological amyloid and tau accumulation. This finding challenges the assumption that the aged APP/PS1 brain represents a functionally inert or irreversible AD state. Instead, it supports the view that even in advanced stages of AD, the brain attempts to mitigate pathology through upregulated endogenous processes (Figure 9, Figure 10A and B). In this context, SCF+G-CSF treatment appears to amplify such mechanisms, including the stimulation of macrophage activity to clear amyloid plaques.^12,85^

Consistent with AD pathology, our analysis also revealed a nearly 15-fold enrichment of complement cascade activation (Figure S5A). Fibrillar amyloid is known to activate both the classical and alternative complement pathways via interactions with C1q and C3, ultimately forming the membrane attack complex (MAC), which damages neuronal membranes and promotes neuroinflammation.^120^ This supports our findings and highlights the role of immune system activation in chronic AD.

Additionally, aged APP/PS1 mice without SCF+G-CSF treatment showed increased enrichment in functions related to hematologic and bone disorders, as well as bacterial, viral, and parasitic infections (Figure S5A). In contrast, SCF+G-CSF-treated APP/PS1 mice exhibited upregulation of genes involved in antimicrobial defense and immune regulation, as described earlier. These findings further support the immunomodulatory and antimicrobial properties of SCF+G-CSF treatment and suggest its potential therapeutic benefit beyond AD.^121–126^ They also reinforce the idea that examining diverse disease processes may provide insights into mechanisms that influence AD progression, resistance, or susceptibility.

## 4. Concluding Remarks

AD is the leading cause of dementia worldwide, with an estimated $350 billion needed to care for individuals aged 65 and older with dementia in the United States in 2024.^127^ Our previous studies have demonstrated that combined SCF+G-CSF treatment reduces AD-related pathologies, including amyloid burden, aggregated tau accumulation, dendritic and synaptic loss, neuroinflammation^11,12^ and cognitive impairments in APP/PS1 mice (Li et al., unpublished observations). Bone marrow-derived macrophages have been shown to effectively phagocytose Aβ^126,127,11,12 11,12^ SCF+G-CSF treatment has shown therapeutic potential across various conditions, including stroke,^121–124^ multiple myeloma,^125^ lymphomas,^126,128^ and recovery of hematopoietic stem/progenitor cells following chemotherapy.^125,129^ Despite these encouraging findings, the molecular mechanisms remain poorly understood. The goal of this study was to use transcriptomic profiling via microarray analysis to explore the mechanisms underlying SCF+G-CSF’s therapeutic effects in aged APP/PS1 mice.

A major strength of the present study lies in its integrated and automated transcriptomic approach. Multiple analytic platforms, including TAC, STRING v12.0, and ShinyGO 0.77, were employed to validate the findings. Additionally, the use of the well-characterized APP/PS1 mouse model, previously shown to respond to SCF+G-CSF treatment ^10,11^, adds credibility to the observed outcomes. Importantly, we examined transcriptomic differences in aged mice, comparing APP/PS1 mice with and without SCF+G-CSF treatment, as well with age-matched WT controls. This allowed us to characterize the AD-related pathological state and its modulation by SCF+G-CSF treatment, highlighting potential molecular pathways and targets involved in AD progression and its mitigation. Thus, this study offers critical insight that can guide future research into the mechanisms and therapeutic potential of SCF+G-CSF treatment in AD.

Nonetheless, the study has limitations. The small sample size is a common constraint in gene chip studies and warrants cautious interpretation. Moreover, while transcriptomic analysis provides valuable insight, confirmatory studies are needed to validate and expand upon the findings. The interpretation of bioinformatic data is also inherently limited by the reliance on existing literature and reference databases, which may introduce bias. Additionally, the analysis tools varied in their capacity to recognize and process gene inputs, which was considered in interpreting the results.

Our study, through the analysis of upregulated and downregulated DEGs, revealed that SCF+G-CSF treatment may act as a metal ion chelator while also stimulating the immune system. It creates a pro-inflammatory environment conducive to microglial activation, enhanced phagocytic activity, amyloid localization, and potentially microglial repair and regeneration (Figures 4, 6, S2, S3), thereby reducing AD pathology in aged APP/PS1 mice.

These mechanisms appear to include, but are not limited to, the upregulation and co-expression of a 6-member gene cluster (S100a8, S100a9, Lcn2, Ngp, Ltf, and Camp) enriched in the spleen and lymphoid tissue (Figure 4). This finding highlights that inflammation, while often viewed as detrimental in AD, may play a necessary role in initiating immune and microglial activation.

Accordingly, our study challenges the conventional neuroinflammatory model of AD by proposing that suppressing inflammation may be an ineffective therapeutic strategy. Instead, we suggest that certain pro-inflammatory responses are essential components of the innate immune system’s ability to mitigate AD pathology. Importantly, the presence of S100A9 at amyloid plaques or the colocalization of S100A8/A9 with amyloid-β should not be misinterpreted as evidence of causality.

We found that IL-17 signaling is highly enriched in SCF+G-CSF-treated APP/PS1 mice compared to vehicle-treated APP/PS1 control mice. We postulate that this IL-17 enrichment is regulated by S100a8, S100a9, and Lcn2 genes (Figures 4, S2), which may facilitate the recruitment of innate and adaptive immune cells into the brain to reduce amyloid burden.^40^ The S100a8/a9 genes also play central roles in oxidative burst pathways, which are critical for degrading intracellular and extracellular debris.^20,23^

Additionally, SCF+G-CSF treatment downregulates EGFR1, a gene known to promote tumorigenesis ^88^. This downregulation may further contribute to the observed reduction in amyloid and tau pathology (Figure 5). Although the link between EGFR1 signaling and AD remains poorly understood, EGFR1 is known to interact with inflammatory mediators such as TNFα and to regulate neural stem cell proliferation in the subventricular zone. ^88^ Its downregulation has been associated with neurodegeneration and a shift toward glial rather than neuronal differentiation, underscoring its importance in maintaining the neural stem/progenitor cell pool. ^88^

Our transcriptomic analysis also revealed that SCF+G-CSF treatment may reduce neuroinflammation in aged APP/PS1 mice by downregulating key cytokine pathways, including IL-7, IL-2, and IL-4, along with MMP16 (Figure 5). This suggests anti-inflammatory effects of SCF+G-CSF, along with the potential to promote extracellular matrix stabilization. Furthermore, the upregulation of Camp and S100a8/a9 may support tissue repair and reduce neurotoxicity^21,27,47^ (Figure 4).

Together with prior findings, our results suggest that SCF+G-CSF may enhance synaptic regeneration and attenuate cognitive decline in AD (Figure 6). These effects may be mediated through metal ion sequestration, which reduces neurotoxicity ^33,34,39,91^ (Figures 4, 6, S3), and through RHO-GTPase-mediated activation of NADPH oxidase pathways, which support actin polymerization and synapse formation. ^95^ (Figures 6, S3). This study also highlights the potential value of exploring cancer-related pathways to better understand the mechanisms underlying AD neuropathology. We found that Cbl, a key cancer-promoting gene, was significantly downregulated by SCF+G-CSF treatment in over 40% of enriched pathways (Figures 5, 6, S3). Notably, 4 out of the top 5 most downregulated genes in SCF+G-CSF-treated APP/PS1 mice (Cbl, Kcnq1ot1, Akap9, Snhg11) are well-established oncogenes (Figure 2B, Table 1). Cancer and neurodegeneration share overlapping features such as chronic inflammation, DNA damage, oxidative stress, and broad metabolic dysregulation.^75,76^

Epidemiological evidence also suggests an inverse relationship between the two conditions: individuals with AD exhibit a lower risk of developing cancer, and cancer survivors show a reduced likelihood of developing AD.^75^

Our findings support the hypothesis that SCF+G-CSF treatment may exert anti-cancer effects, as reflected in the consistent downregulation of cancer-promoting genes, particularly Cbl. Other oncogenes, including Kcnq1ot1 and Akap9, also showed decreased expression following SCF+G-CSF treatment. These results suggest that targeting overlapping molecular mechanisms between cancer and neurodegeneration could offer a novel therapeutic angle in treating AD and warrant further investigation.

Overall, our findings highlight that reducing AD pathology by blocking neuroinflammation *alone* may impair essential microglial and innate immune functions. While many current therapeutic approaches aim to suppress neuroinflammation or dampen immune responses in AD ^2,4,5^, our results suggest that microglial activation is enhanced under pro-inflammatory conditions. Our study challenges the prevailing neuroinflammatory model of AD and suggests that effective treatment strategies for AD should not only *promote*, but also *tightly regulate,* a permissive inflammatory environment to optimize innate and adaptive immune functions necessary for amyloid clearance.

Figure 11 provides an overview of the complex and diverse mechanisms through which we propose SCF+G-CSF treatment reduces Alzheimer’s disease neuropathology. In addition to the classical hallmarks of AD, our study reinforces the significance of metal ion dys-homeostasis, immune dysregulation, and overlapping features with cancer pathology. These insights reveal novel therapeutic targets and mechanisms by which SCF+G-CSF treatment may reduce AD pathology in aged APP/PS1 mice. Moving forward, AD research and treatment development should more fully integrate these multifaceted mechanisms to more effectively address the global burden of Alzheimer’s disease.

**Figure 11.**
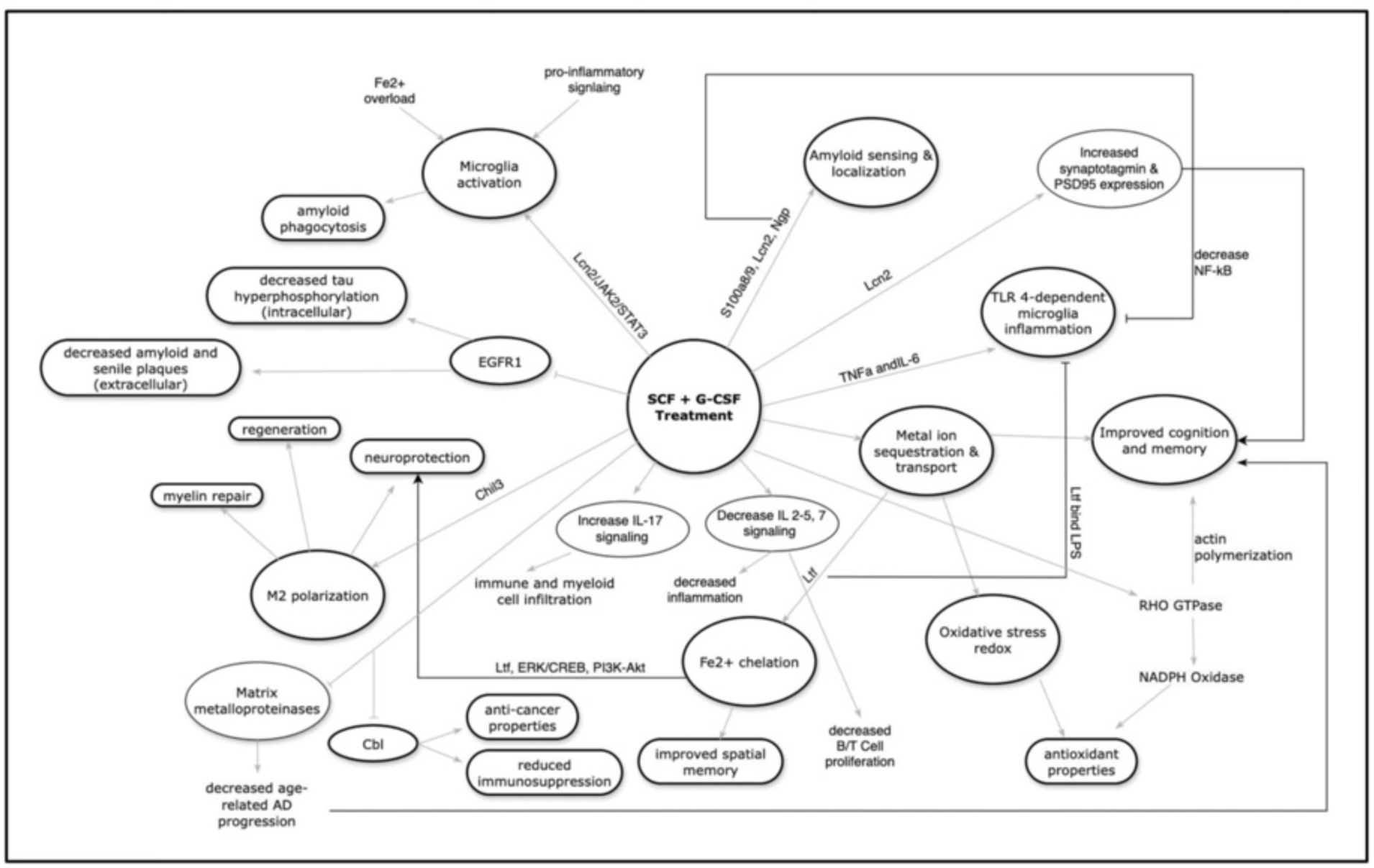
Proposed mechanism of SCF+G-CSF treatment in aged APP/PS1 mice.

## Supporting information

Suppemental figures and table

## Author Contributions

Conceptualization, L.-R.Z.; methodology, L.-R.Z.; formal analysis, A. A.; investigation, A. A., B.L., S.M.; resources, L.-R.Z.; data curation, A. A., B.L., S.M.; writing—first draft preparation, A. A.; writing—review and editing, L.-R.Z.; visualization, A. A., R.G.; supervision, L.-R.Z.; project administration and funding acquisition, L.-R.Z.

## Availability of data and materials

The datasets generated during the current study are available from the corresponding author on reasonable request.

## Competing Interests

The authors declare no conflict of interest.

## Acknowledgement

This study was supported by the Frank’s Foundation.

## Notes

### Competing Interest Statement

The authors have declared no competing interest.

### Summary of Updates

Alzheimers Disease is a neurodegenerative disease characterized by amyloid plaque deposition, tau hyperphosphorylation, neuroinflammation, and cognitive decline. Our previous studies showed that combined treatment with stem cell factor, SCF, and granulocyte colony stimulating factor, GCSF, reduces Alzheimers Disease pathology in APPPS1 mice. This study aimed to explore the molecular mechanism underlying the therapeutic effects using transcriptomic analysis. Aged APPPS1 mice received daily subcutaneous injections of SCF and GCSF or vehicle for 12 days. RNA was extracted from brain tissue on day 13 for gene chip analysis. Age matched wild type, WT, mice served as controls. Data were analyzed using TAC, STRING v12, Reactome, and ShinyGO. A total of 45037 differentially expressed genes, DEGs, were detected. Twenty seven DEGs met a more than two fold threshold in SCF and GCSF treated versus vehicle treated APPPS1 mice; 89 DEGs met this threshold in APPPS1 versus WT mice. SCF and GCSF treatment upregulated six immune related genes, S100a8, S100a9, Ngp, Lcn2, Ltf, and Camp, associated with amyloid clearance, immune cell recruitment, and repair. Pathway analysis showed downregulation of IL2, IL4, IL7, and EGFR1, and upregulation of IL17 signaling, suggesting modulation of both innate and adaptive immunity. Notably, SCF and GCSF downregulated several oncogenes, including Cbl, Akap9, Kcnq1ot1, and Snhg11, highlighting an overlap between cancer and Alzheimers Disease related pathways. SCF and GCSF also promoted NADPH oxidase activation via Rho GTPases and showed more than 400 fold enrichment in metal ion sequestration, indicating potential metal chelation effects. These findings suggest that SCF and GCSF treatment modifies immune and metabolic pathways, reduces Alzheimers Disease pathology, and highlights new therapeutic targets involving inflammation, metal homeostasis, and oncogenic signaling.

## References

1. Long JM, Holtzman DM. Alzheimer disease: An update on pathobiology and treatment strategies. Cell. 2019;179(2):312–339. https://dx.doi.org/10.1016/j.cell.2019.09.001. doi: 10.1016/j.cell.2019.09.001.

2. Rajesh Y, Kanneganti T. Innate immune cell death in neuroinflammation and alzheimer’s disease. Cells (Basel, Switzerland). 2022;11(12):1885. https://search.proquest.com/docview/2679686848. doi: 10.3390/cells11121885.

3. 2025 alzheimer’s disease facts and figuresAlzheimer’s & dementia. 2025;21(4):n/a. https://onlinelibrary.wiley.com/doi/abs/10.1002%2Falz.70235. doi: 10.1002/alz.70235.

4. Wong W. Economic burden of alzheimer disease and managed care considerations. The American journal of managed care. 2020;26(8 Suppl):S177–S183. https://www.ncbi.nlm.nih.gov/pubmed/32840331. doi: 10.37765/ajmc.2020.88482.

5. Potter H, Woodcock JH, Boyd TD, et al. Safety and efficacy of sargramostim (GM-CSF) in the treatment of alzheimer’s disease. Alzheimer’s & dementia : translational research & clinical interventions. 2021;7(1):e12158–n/a. https://onlinelibrary.wiley.com/doi/abs/10.1002/trc2.12158. doi: 10.1002/trc2.12158.

6. Lampron A, Gosselin D, Rivest S. Targeting the hematopoietic system for the treatment of alzheimer’s disease. Brain, behavior, and immunity. 2011;25:S71–S79. https://www.clinicalkey.es/playcontent/1-s2.0-S088915911000591X. doi:10.1016/j.bbi.2010.12.018.

7. Groarke EM, Young NS. Aging and hematopoiesis. Clinics in Geriatric Medicine. 2019;35(3):285–293. doi: 10.1016/j.cger.2019.03.001.

8. Yan P, Kim K, Xiao Q, et al. Peripheral monocyte–derived cells counter amyloid plaque pathogenesis in a mouse model of alzheimer’s disease. The Journal of clinical investigation. 2022;132(11). https://www.ncbi.nlm.nih.gov/pubmed/35511433. doi: 10.1172/JCI152565.

9. Bouzid H, Belk JA, Jan M, et al. Clonal hematopoiesis is associated with protection from alzheimer’s disease. Nature medicine. 2023;29(7):1662–1670. https://www.ncbi.nlm.nih.gov/pubmed/37322115. doi: 10.1038/s41591-023-02397-2.

10. Wyss-Coray T, Ray S, Britschgi M, et al. Classification and prediction of clinical alzheimer’s diagnosis based on plasma signaling proteins. Nature medicine. 2007;13(11):1359–1362. http://dx.doi.org/10.1038/nm1653. doi: 10.1038/nm1653.

11. Guo X, Liu Y, Morgan D, Zhao L. Reparative effects of stem cell factor and granulocyte colony-stimulating factor in aged APP/PS1 mice. Aging and disease. 2020;11(6):1423–1443. https://www.ncbi.nlm.nih.gov/pubmed/33269098. doi: 10.14336/AD.2020.0201.

12. Li B, Gonzalez-Toledo ME, Piao C, Gu A, Kelley RE, Zhao L. Stem cell factor and granulocyte colony-stimulating factor reduce β-amyloid deposits in the brains of APP/PS1 transgenic mice. Alzheimer’s research & therapy. 2011;3(2):8. https://www.ncbi.nlm.nih.gov/pubmed/21406112. doi: 10.1186/alzrt67.

13. ThermoFisher Scientific. Transcriptome analysis console (RRID:SCR_016519).

14. Szklarczyk D, Kirsch R, Koutrouli M, et al. The STRING database in 2023: Protein– Protein association networks and functional enrichment analyses for any sequenced genome of interest. Nucleic acids research. 2023;51(D1):D638–D646. https://www.ncbi.nlm.nih.gov/pubmed/36370105. doi: 10.1093/nar/gkac1000.

15. Ge SX, Jung D, Yao R. ShinyGO: A graphical gene-set enrichment tool for animals and plants. Bioinformatics. 2020;36(8):2628–2629. https://www.ncbi.nlm.nih.gov/pubmed/31882993. doi: 10.1093/bioinformatics/btz931.

16. Griss J, Viteri G, Sidiropoulos K, Nguyen V, Fabregat A, Hermjakob H. ReactomeGSA - efficient multi-omics comparative pathway analysis. Molecular & cellular proteomics. 2020;19(12):2115–2125. https://dx.doi.org/10.1074/mcp.TIR120.002155. doi: 10.1074/mcp.TIR120.002155.

17. Radde R, Bolmont T, Kaeser SA, et al. Abeta42-driven cerebral amyloidosis in transgenic mice reveals early and robust pathology. EMBO reports. 2006;7(9):940–946. https://www.ncbi.nlm.nih.gov/pubmed/16906128.

18. Elkon R, Agami R. Characterization of noncoding regulatory DNA in the human genome. Nat Biotechnol. 2017;35(8):732–746. https://link.springer.com/article/10.1038/nbt.3863. doi: 10.1038/nbt.3863.

19. Statello L, Guo C, Chen L, Huarte M. Gene regulation by long non-coding RNAs and its biological functions. Nat Rev Mol Cell Biol. 2021;22(2):96–118. https://link.springer.com/article/10.1038/s41580-020-00315-9. doi: 10.1038/s41580-020-00315-9.

20. Markowitz J, Carson WE. Review of S100A9 biology and its role in cancer. Biochimica et biophysica acta. 2013;1835(1):100–109. https://dx.doi.org/10.1016/j.bbcan.2012.10.003. doi: 10.1016/j.bbcan.2012.10.003.

21. Pruenster M, Vogl T, Roth J, Sperandio M. S100A8/A9: From basic science to clinical application. Pharmacology & Therapeutics. 2016;167:120–131. https://www.ncbi.nlm.nih.gov/pubmed/27492899. doi: 10.1016/j.pharmthera.2016.07.015.

22. Cristóvão JS, Gomes CM. S100 proteins in alzheimer’s disease. Frontiers in neuroscience. 2019;13:463. https://www.ncbi.nlm.nih.gov/pubmed/31156365. doi: 10.3389/fnins.2019.00463.

23. Lim SY, Raftery MJ, Goyette J, Hsu K, Geczy CL. Oxidative modifications of S100 proteins: Functional regulation by redox. Journal of Leukocyte Biology. 2009;86(3):577–587. http://www.jleukbio.org/content/86/3/577.abstract. doi: 10.1189/jlb.1008608.

24. Malm TM, Koistinaho M, Pärepalo M, et al. Bone-marrow-derived cells contribute to the recruitment of microglial cells in response to β-amyloid deposition in APP/PS1 double transgenic alzheimer mice. Neurobiology of disease. 2005;18(1):134–142. https://dx.doi.org/10.1016/j.nbd.2004.09.009. doi: 10.1016/j.nbd.2004.09.009.

25. Wolf M, Joseph R, Austermann J, et al. S100A8/S100A9 integrates F-actin and microtubule dynamics to prevent uncontrolled extravasation of leukocytes. Biomedicines. 2023;11(3):835. https://www.ncbi.nlm.nih.gov/pubmed/36979814. doi:10.3390/biomedicines11030835.

26. Ma L, Sun P, Zhang J, Zhang Q, Yao S. Proinflammatory effects of S100A8/A9 via TLR4 and RAGE signaling pathways in BV-2 microglial cells. International Journal of Molecular Medicine. 2017;40(1):31–38. https://www.ncbi.nlm.nih.gov/pubmed/28498464. doi: 10.3892/ijmm.2017.2987.

27. Willis EF, MacDonald KPA, Nguyen QH, et al. Repopulating microglia promote brain repair in an IL-6-dependent manner. Cell. 2020;180(5):833–846.e16. https://dx.doi.org/10.1016/j.cell.2020.02.013. doi: 10.1016/j.cell.2020.02.013.

28. Hagmeyer S, Romão MA, Cristóvão JS, et al. Distribution and relative abundance of S100 proteins in the brain of the APP23 alzheimer’s disease model mice. Frontiers in neuroscience. 2019;13:640. https://www.ncbi.nlm.nih.gov/pubmed/31281238. doi:10.3389/fnins.2019.00640.

29. Dekens DW, Naudé PJW, Engelborghs S, et al. Neutrophil gelatinase-associated lipocalin and its receptors in alzheimer’s disease (AD) brain regions: Differential findings in AD with and without depression. Journal of Alzheimer’s disease. 2017;55(2):763–776. https://www.ncbi.nlm.nih.gov/pubmed/27716662. doi: 10.3233/JAD-160330.

30. Jaberi SA, Cohen A, D’Souza C, et al. Lipocalin-2: Structure, function, distribution and role in metabolic disorders. Biomedicine & Pharmacotherapy. 2021;142:112002. https://dx.doi.org/10.1016/j.biopha.2021.112002. doi: 10.1016/j.biopha.2021.112002.

31. Kang SS, Ren Y, Liu C, et al. Lipocalin-2 protects the brain during inflammatory conditions. Molecular Psychiatry. 2018;23(2):344–350. https://www.ncbi.nlm.nih.gov/pubmed/28070126. doi: 10.1038/mp.2016.243.

32. Yang H, Wang X, Li S, Liu Y, Akbar R, Fan G. Lipocalin family proteins and their diverse roles in cardiovascular disease. Pharmacology & therapeutics (Oxford). 2023;244:108385. https://dx.doi.org/10.1016/j.pharmthera.2023.108385. doi:10.1016/j.pharmthera.2023.108385.

33. Mesquita SD, Ferreira AC, Falcao AM, et al. Lipocalin 2 modulates the cellular response to amyloid beta. Cell Death & Differentiation. 2014;21(10):1588–1599. https://www.ncbi.nlm.nih.gov/pubmed/24853299. doi: 10.1038/cdd.2014.68.

34. Ferreira AC, Dá Mesquita S, Sousa JC, et al. From the periphery to the brain: Lipocalin-2, a friend or foe? Progress in Neurobiology. 2015;131:120–136. https://www.ncbi.nlm.nih.gov/pubmed/26159707. doi: 10.1016/j.pneurobio.2015.06.005.

35. Bathini P, Dupanloup I, Zenaro E, et al. Systemic inflammation causes microglial dysfunction with a vascular AD phenotype. Brain, behavior, & immunity. Health. 2023;28:100568. https://dx.doi.org/10.1016/j.bbih.2022.100568. doi:10.1016/j.bbih.2022.100568.

36. Llorens F, Hermann P, Villar-Piqué A, et al. Cerebrospinal fluid lipocalin 2 as a novel biomarker for the differential diagnosis of vascular dementia. Nature Communications. 2020;11(1):619. https://www.ncbi.nlm.nih.gov/pubmed/32001681. doi: 10.1038/s41467-020-14373-2.

37. Berger T, Togawa A, Duncan GS, et al. Lipocalin 2-deficient mice exhibit increased sensitivity to escherichia coli infection but not to ischemia-reperfusion injury. Proceedings of the National Academy of Sciences - PNAS. 2006;103(6):1834–1839. https://www.jstor.org/stable/30048465. doi: 10.1073/pnas.0510847103.

38. Aderem A, Flo TH, Smith KD, et al. Lipocalin 2 mediates an innate immune response to bacterial infection by sequestrating iron. Nature. 2004;432(7019):917–921. https://dx.doi.org/10.1038/nature03104. doi: 10.1038/nature03104.

39. Cline EN, Bicca MA, Viola KL, Klein WL. The amyloid-β oligomer hypothesis: Beginning of the third decade. Journal of Alzheimer’s disease. 2018;64(s1):S567–S610. https://www.ncbi.nlm.nih.gov/pubmed/29843241. doi: 10.3233/JAD-179941.

40. Adler O, Zait Y, Cohen N, et al. Reciprocal interactions between innate immune cells and astrocytes facilitate neuroinflammation and brain metastasis via lipocalin-2. Nature cancer. 2023;4(3):401–418. https://www.ncbi.nlm.nih.gov/pubmed/36797502. doi: 10.1038/s43018-023-00519-w.

41. Moscinski LNC, Hill B. Molecular cloning of a novel myeloid granule protein. Journal of cellular biochemistry. 1995;59(4):431–442. https://api.istex.fr/ark:/67375/WNG-TP97QZX2-8/fulltext.pdf. doi: 10.1002/jcb.240590404.

42. Boutté AM, Friedman DB, Bogyo M, Min Y, Yang L, Lin PC. Identification of a myeloid-derived suppressor cell cystatin-like protein that inhibits metastasis. The FASEB journal. 2011;25(8):2626–2637. https://onlinelibrary.wiley.com/doi/abs/10.1096%2Ffj.10-180604. doi: 10.1096/fj.10-180604.

43. Mutua V, Gershwin LJ. A review of neutrophil extracellular traps (NETs) in disease: Potential anti-NETs therapeutics. Clinic Rev Allerg Immunol. 2021;61(2):194–211. https://link.springer.com/article/10.1007/s12016-020-08804-7. doi: 10.1007/s12016-020-08804-7.

44. Vorobjeva NV, Chernyak BV. NETosis: Molecular mechanisms, role in physiology and pathology. Biochemistry Moscow. 2020;85(10):1178–1190. https://link.springer.com/article/10.1134/S0006297920100065. doi:10.1134/s0006297920100065.

45. Pietronigro EC, Della Bianca V, Zenaro E, Constantin G. NETosis in alzheimer’s disease. Frontiers in immunology. 2017;8:211. https://www.ncbi.nlm.nih.gov/pubmed/28303140. doi: 10.3389/fimmu.2017.00211.

46. Smyth LCD, Murray HC, Hill M, et al. Neutrophil-vascular interactions drive myeloperoxidase accumulation in the brain in alzheimer’s disease. Acta neuropathologica communications. 2022;10(1):38. https://www.ncbi.nlm.nih.gov/pubmed/35331340. doi: 10.1186/s40478-022-01347-2.

47. Orihuela R, McPherson CA, Harry GJ. Microglial M1/M2 polarization and metabolic states. British Journal of Pharmacology. 2016;173(4):649–665. https://onlinelibrary.wiley.com/doi/abs/10.1111%2Fbph.13139. doi: 10.1111/bph.13139.

48. Hao L, Shan Q, Wei J, Ma F, Sun P. Lactoferrin: Major physiological functions and applications. Current Protein & Peptide Science. 2019;20(2):139–144. http://www.eurekaselect.com/openurl/content.php?genre=article&issn=1389-2037&volume=20&issue=2&spage=139. doi: 10.2174/1389203719666180514150921.

49. Li Y, Guo C. A review on lactoferrin and central nervous system diseases. Cells. 2021;10(7):1810. https://search.proquest.com/docview/2554473674. doi: 10.3390/cells10071810.

50. Khan AI, Liu J, Dutta P. Bayesian inference for parameter estimation in lactoferrin-mediated iron transport across blood-brain barrier. Biochimica et biophysica acta. G, General subjects/Biochimica et biophysica acta. General subjects (Online). 2020;1864(3):129459. https://dx.doi.org/10.1016/j.bbagen.2019.129459. doi:10.1016/j.bbagen.2019.129459.

51. Guo C, Yang Z, Zhang S, et al. Intranasal lactoferrin enhances α-secretase-dependent amyloid precursor protein processing via the ERK1/2-CREB and HIF-1α pathways in an alzheimer’s disease mouse model. Neuropsychopharmacology. 2017;42(13):2504–2515. https://www.ncbi.nlm.nih.gov/pubmed/28079060. doi:10.1038/npp.2017.8.

52. Bhusal A, Nam Y, Seo D, Lee W, Suk K. Cathelicidin-related antimicrobial peptide negatively regulates bacterial endotoxin-induced glial activation. Cells. 2022;11(23):3886. https://www.ncbi.nlm.nih.gov/pubmed/36497142. doi:10.3390/cells11233886.

53. Guo T, Zhang D, Zeng Y, Huang TY, Xu H, Zhao Y. Molecular and cellular mechanisms underlying the pathogenesis of alzheimer’s disease. Molecular Neurodegeneration. 2020;15(1):1–40. https://www.proquest.com/docview/2424809085/abstract/. doi: 10.1186/s13024-020-00391-7.

54. Steel AJ, Eslick GD. Herpes viruses increase the risk of alzheimer’s disease: A meta-analysis. Journal of Alzheimer’s disease. 2015;47(2):351–364. https://www.ncbi.nlm.nih.gov/pubmed/26401558. doi: 10.3233/JAD-140822.

55. Gombart AF, Borregaard N, Koeffler HP. Human cathelicidin antimicrobial peptide (CAMP) gene is a direct target of the vitamin D receptor and is strongly up-regulated in myeloid cells by 1,25-dihydroxyvitamin D3. The FASEB Journal. 2005;19(9):1067–1077. https://onlinelibrary.wiley.com/doi/abs/10.1096%2Ffj.04-3284com. doi: 10.1096/fj.04-3284com.

56. Lowry MB, Guo C, Zhang Y, et al. A mouse model for vitamin D-induced human cathelicidin antimicrobial peptide gene expression. The Journal of steroid biochemistry and molecular biology. 2020;198:105552. https://dx.doi.org/10.1016/j.jsbmb.2019.105552. doi: 10.1016/j.jsbmb.2019.105552.

57. Wang G, Chen Z, Tian P, et al. Wound healing mechanism of antimicrobial peptide cathelicidin-DM. Frontiers in bioengineering and biotechnology. 2022;10:977159. https://search.proquest.com/docview/2740502623. doi: 10.3389/fbioe.2022.977159.

58. Mangoni ML, McDermott AM, Zasloff M. Antimicrobial peptides and wound healing: Biological and therapeutic considerations. Experimental Dermatology. 2016;25(3):167–173. https://api.istex.fr/ark:/67375/WNG-JLCM4WHS-L/fulltext.pdf. doi:10.1111/exd.12929.

59. Karadurmus D, Rial D, De Backer J, Communi D, de Kerchove d’Exaerde A, Schiffmann SN. GPRIN3 controls neuronal excitability, morphology, and striatal-dependent behaviors in the indirect pathway of the striatum. The Journal of neuroscience. 2019;39(38):7513–7528. https://www.ncbi.nlm.nih.gov/pubmed/31363062. doi: 10.1523/JNEUROSCI.2454-18.2019.

60. Yasumasa Mototani, Tadashi Okamura, Motohito Goto, et al. Role of G protein-regulated inducer of neurite outgrowth 3 (GRIN3) in β-arrestin 2-akt signaling and dopaminergic behaviors. Pflügers Archiv - European Journal of Physiology. 2018;470(6):937–947. doi: 10.1007/s00424-018-2124-1.

61. Tang D, Luo Y, Jiang Y, et al. LncRNA KCNQ1OT1 activated by c-myc promotes cell proliferation via interacting with FUS to stabilize MAP3K1 in acute promyelocytic leukemia. Cell death & disease. 2021;12(9):795. https://www.ncbi.nlm.nih.gov/pubmed/34404765. doi: 10.1038/s41419-021-04080-1.

62. Cagle P, Qi Q, Niture S, Kumar D. KCNQ1OT1: An oncogenic long noncoding RNA. Biomolecules. 2021;11(11):1602. https://www.ncbi.nlm.nih.gov/pubmed/34827600. doi:10.3390/biom11111602.

63. Ghiam S, Eslahchi C, Shahpasand K, Habibi-Rezaei M, Gharaghani S. Exploring the role of non-coding RNAs as potential candidate biomarkers in the cross-talk between diabetes mellitus and alzheimer’s disease. Frontiers in aging neuroscience. 2022;14:955461. https://www.ncbi.nlm.nih.gov/pubmed/36092798. doi:10.3389/fnagi.2022.955461.

64. Jaberi KR, Alamdari-Palangi V, Jaberi AR, et al. The regulation, functions, and signaling of miR-153 in neurological disorders, and its potential as a biomarker and therapeutic target. Current molecular medicine. 2023;23(9):863–875. http://www.eurekaselect.com/openurl/content.php?genre=article&issn=1875-5666&volume=23&issue=9&spage=863. doi: 10.2174/1566524023666220817145638.

65. Huang X, Tan J, Li Y, et al. Expression of LncRNA KCNQ1Ot1 in diabetic nephropathy and its correlation with MEK/ERK signaling pathway. American journal of translational research. 2022;14(3):1796–1806. https://www.ncbi.nlm.nih.gov/pubmed/35422925.

66. Rapetti-Mauss R, Bustos V, Thomas W, et al. Bidirectional KCNQ1. Proceedings of the National Academy of Sciences - PNAS. 2017;114(16):4159–4164. https://www.jstor.org/stable/26480423. doi: 10.1073/pnas.1702913114.

67. Herter JM, Grabie N, Cullere X, et al. AKAP9 regulates activation-induced retention of T lymphocytes at sites of inflammation. Nature communications. 2015;6(1):10182. https://www.ncbi.nlm.nih.gov/pubmed/26680259. doi: 10.1038/ncomms10182.

68. Ikezu T, Chen C, DeLeo AM, et al. Tau phosphorylation is impacted by rare AKAP9 mutations associated with alzheimer disease in african americans. J Neuroimmune Pharmacol. 2018;13(2):254–264. https://link.springer.com/article/10.1007/s11481-018-9781-x. doi: 10.1007/s11481-018-9781-x.

69. Wu S, Shen D, Zhao L. AKAP9 upregulation predicts unfavorable prognosis in pediatric acute myeloid leukemia and promotes stemness properties via the wnt/β-catenin pathway. Cancer management and research. 2022;14:157–167. https://www.ncbi.nlm.nih.gov/pubmed/35046723. doi: 10.2147/CMAR.S343033.

70. Reddy JS, Jin J, Lincoln SJ, et al. Transcript levels in plasma contribute substantial predictive value as potential alzheimer’s disease biomarkers in african americans. EBioMedicine. 2022;78:103929. https://dx.doi.org/10.1016/j.ebiom.2022.103929. doi:10.1016/j.ebiom.2022.103929.

71. You Y, Hersh SW, Aslebagh R, et al. Alzheimer’s disease associated AKAP9 I2558M mutation alters posttranslational modification and interactome of tau and cellular functions in CRISPR-edited human neuronal cells. Aging Cell. 2022;21(6):e13617–n/a. https://onlinelibrary.wiley.com/doi/abs/10.1111%2Facel.13617. doi: 10.1111/acel.13617.

72. Kales SC, Ryan PE, Nau MM, Lipkowitz S. Cbl and human myeloid neoplasms: The cbl oncogene comes of age. Cancer Research. 2010;70(12):4789–4794. https://www.ncbi.nlm.nih.gov/pubmed/20501843. doi: 10.1158/0008-5472.can-10-0610.

73. Sattler M, Pride YB, Quinnan LR, et al. Differential expression and signaling of CBL and CBL-B in BCR/ABL transformed cells. Oncogene. 2002;21(9):1423–1433. https://www.ncbi.nlm.nih.gov/pubmed/11857085. doi: 10.1038/sj.onc.1205202.

74. Lyle CL, Belghasem M, Chitalia VC. C-cbl: An important regulator and a target in angiogenesis and tumorigenesis. Cells. 2019;8(5):498. https://www.ncbi.nlm.nih.gov/pubmed/31126146. doi: 10.3390/cells8050498.

75. Driver JA, Beiser A, Au R, et al. Inverse association between cancer and alzheimer’s disease: Results from the framingham heart study. BMJ. 2012;344(mar12 1):1442. https://dx.doi.org/10.1136/bmj.e1442. doi: 10.1136/bmj.e1442.

76. Houck AL, Seddighi S, Driver JA. At the crossroads between neurodegeneration and cancer: A review of overlapping biology and its implications. Current aging science. 2018;11(2):77–89. http://www.eurekaselect.com/openurl/content.php?genre=article&issn=1874-6098&volume=11&issue=2&spage=77. doi: 10.2174/1874609811666180223154436.

77. Wu Q, Ma J, Wei J, Meng W, Wang Y, Shi M. lncRNA SNHG11 promotes gastric cancer progression by activating the wnt/β-catenin pathway and oncogenic autophagy. Molecular therapy. 2021;29(3):1258–1278. https://dx.doi.org/10.1016/j.ymthe.2020.10.011. doi: 10.1016/j.ymthe.2020.10.011.

78. Rapetti-Mauss R, Berenguier C, Allegrini B, Soriani O. Interplay between ion channels and the wnt/β-catenin signaling pathway in cancers. Frontiers in pharmacology. 2020;11:525020. https://search.proquest.com/docview/2455836623. doi:10.3389/fphar.2020.525020.

79. Papakonstantinou E, 1, 2, et al. NOTCH3 and CADASIL syndrome: A genetic and structural overview . .

80. Sierra C, Sabariego-Navarro M, Fernández-Blanco Á, et al. The lncRNA Snhg11, a new candidate contributing to neurogenesis, plasticity, and memory deficits in down syndrome. Molecular psychiatry. 2024. https://www.ncbi.nlm.nih.gov/pubmed/38409595. doi: 10.1038/s41380-024-02440-9.

81. Xu L, Huan L, Guo T, et al. LncRNA SNHG11 facilitates tumor metastasis by interacting with and stabilizing HIF-1α. Oncogene. 2020;39(46):7005–7018. https://www.ncbi.nlm.nih.gov/pubmed/33060856. doi: 10.1038/s41388-020-01512-8.

82. Yu Z, Chen D, Tan C, et al. Physiological clearance of aβ by spleen and splenectomy aggravates alzheimer-type pathogenesis. Aging Cell. 2022;21(1):e13533– n/a. https://onlinelibrary.wiley.com/doi/abs/10.1111%2Facel.13533. doi:10.1111/acel.13533.

83. Ye X, Chen J, Pan J, et al. Interleukin-17 promotes the infiltration of CD8+ T cells into the brain in a mouse model for alzheimer’s disease. Immunological investigations. 2023;52(2):135–153. https://www.tandfonline.com/doi/abs/10.1080/08820139.2022.2136525. doi:10.1080/08820139.2022.2136525.

84. Unger MS, Li E, Scharnagl L, et al. CD8+ T-cells infiltrate alzheimer’s disease brains and regulate neuronal- and synapse-related gene expression in APP-PS1 transgenic mice. Brain, behavior, and immunity. 2020;89:67–86. https://dx.doi.org/10.1016/j.bbi.2020.05.070. doi: 10.1016/j.bbi.2020.05.070.

85. Gardner RS, Kyle M, Hughes K, Zhao L. Single-cell RNA sequencing reveals immunomodulatory effects of stem cell factor and granulocyte colony-stimulating factor treatment in the brains of aged APP/PS1 mice. Biomolecules (Basel, Switzerland). 2024;14(7):827. https://www.ncbi.nlm.nih.gov/pubmed/39062541. doi:10.3390/biom14070827.

86. Endo, K., Takeshita, T., Kasai, H., Sasaki, Y., Tanaka, N., Asao, H.,… & Sugamura, K. STAM2, a new member of the STAM family, binding to the janus kinases . .

87. Fromm JA, Johnson SAS, Johnson DL. Epidermal growth factor receptor 1 (EGFR1) and its variant EGFRvIII regulate TATA-binding protein expression through distinct pathways. Molecular and Cellular Biology. 2008;28(20):6483–6495. http://mcb.asm.org/content/28/20/6483.abstract. doi: 10.1128/MCB.00288-08.

88. Romano R, Bucci C. Role of EGFR in the nervous system. Cells. 2020;9(8):1887. https://www.ncbi.nlm.nih.gov/pubmed/32806510. doi: 10.3390/cells9081887.

89. Tavassoly O, Tavassoly I. EGFR aggregation in the brain. ACS chemical neuroscience. 2021;12(11):1833–1834. https://dx.doi.org/10.1021/acschemneuro.1c00264. doi:10.1021/acschemneuro.1c00264.

90. Jayaswamy PK, Vijaykrishnaraj M, Patil P, Alexander LM, Kellarai A, Shetty P. Implicative role of epidermal growth factor receptor and its associated signaling partners in the pathogenesis of alzheimer’s disease. Ageing research reviews. 2023;83:101791. https://dx.doi.org/10.1016/j.arr.2022.101791. doi: 10.1016/j.arr.2022.101791.

91. Wang X, Tan M, Yu J, Tan L. Matrix metalloproteinases and their multiple roles in alzheimer’s disease. BioMed research international. 2014;2014:908636–8. https://dx.doi.org/10.1155/2014/908636. doi: 10.1155/2014/908636.

92. Tjalkens RB, Popichak KA, Kirkley KA. Inflammatory activation of microglia and astrocytes in manganese neurotoxicity. In: Advances in neurobiology. Vol 18. Cham: Springer International Publishing; 2017:159–181. http://link.springer.com/10.1007/978-3-319-60189-2_8. 10.1007/978-3-319-60189-2_8.

93. Chen L, Fan Y, Zhao L, Zhang Q, Wang Z. The metal ion hypothesis of alzheimer’s disease and the anti-neuroinflammatory effect of metal chelators. Bioorganic chemistry. 2023;131:106301. https://dx.doi.org/10.1016/j.bioorg.2022.106301. doi:10.1016/j.bioorg.2022.106301.

94. Liu F, Zhang Z, Zhang L, et al. Effect of metal ions on alzheimer’s disease. Brain and behavior. 2022;12(3):e2527–n/a. https://onlinelibrary.wiley.com/doi/abs/10.1002/brb3.2527. doi: 10.1002/brb3.2527.

95. Aguilar BJ, Zhu Y, Lu Q. Rho GTPases as therapeutic targets in alzheimer’s disease. Alzheimer’s Research & Therapy. 2017;9(1):97. https://www.ncbi.nlm.nih.gov/pubmed/29246246. doi: 10.1186/s13195-017-0320-4.

96. Ma J, Jiang T, Tan L, Yu J. TYROBP in alzheimer’s disease. Mol Neurobiol. 2015;51(2):820–826. https://link.springer.com/article/10.1007/s12035-014-8811-9. doi:10.1007/s12035-014-8811-9.

97. Li R, Lv Z, Li Y, Li W, Hao Y. Effects of TYROBP deficiency on neuroinflammation of a alzheimer’s disease mouse model carrying a PSEN1 p.G378E mutation. Chinese medical sciences journal. 2022;37(4):320–330. https://dx.doi.org/10.24920/004059. doi:10.24920/004059.

98. Kos J, Nanut MP, Prunk M, Sabotič J, Dautović E, Jewett A. Cystatin F as a regulator of immune cell cytotoxicity. Cancer Immunol Immunother. 2018;67(12):1931– 1938. https://link.springer.com/article/10.1007/s00262-018-2165-5. doi:10.1007/s00262-018-2165-5.

99. Daniels MJD, Lefevre L, Szymkowiak S, et al. Cystatin F (Cst7) drives sex-dependent changes in microglia in an amyloid-driven model of alzheimer’s disease. eLife. 2023;12. https://www.ncbi.nlm.nih.gov/pubmed/38085657. doi:10.7554/eLife.85279.

100. Li Q, Li B, Liu L, et al. Monocytes release cystatin F dimer to associate with aβ and aggravate amyloid pathology and cognitive deficits in alzheimer’s disease. Journal of neuroinflammation. 2024;21(1):125. https://www.ncbi.nlm.nih.gov/pubmed/38730470. doi: 10.1186/s12974-024-03119-2.

101. Ma J, Tanaka KF, Shimizu T, et al. Microglial cystatin F expression is a sensitive indicator for ongoing demyelination with concurrent remyelination. Journal of neuroscience research. 2011;89(5):639–649. https://api.istex.fr/ark:/67375/WNG-D1RJ5JTS-7/fulltext.pdf. doi: 10.1002/jnr.22567.

102. Wang S, Sudan R, Peng V, et al. TREM2 drives microglia response to amyloid-β via SYK-dependent and -independent pathways. Cell. 2022;185(22):4153–4169.e19. https://www.ncbi.nlm.nih.gov/pubmed/36306735. doi: 10.1016/j.cell.2022.09.033.

103. Brown GD, Taylor PR, Reid DM, et al. Dectin-1 is a major beta-glucan receptor on macrophages. The Journal of experimental medicine. 2002;196(3):407–412. https://www.ncbi.nlm.nih.gov/pubmed/12163569. doi: 10.1084/jem.20020470.

104. Yao H, Coppola K, Schweig JE, Crawford F, Mullan M, Paris D. Distinct signaling pathways regulate TREM2 phagocytic and NFκB antagonistic activities. Frontiers in cellular neuroscience. 2019;13:457. https://www.ncbi.nlm.nih.gov/pubmed/31649511. doi: 10.3389/fncel.2019.00457.

105. Wang J, Li X, Wang K, et al. CLEC7A regulates M2 macrophages to suppress the immune microenvironment and implies poorer prognosis of glioma. Frontiers in immunology. 2024;15:1361351. https://www.proquest.com/docview/3065979542/abstract/. doi:10.3389/fimmu.2024.1361351.

106. Lopes KdP, Yu L, Shen X, et al. Associations of cortical SPP1 and ITGAX with cognition and common neuropathologies in older adults. Alzheimer’s & dementia. 2024;20(1):525–537. https://onlinelibrary.wiley.com/doi/abs/10.1002%2Falz.13474. doi: 10.1002/alz.13474.

107. Cudaback E, Yang Y, Montine TJ, Keene CD. APOE genotype-dependent modulation of astrocyte chemokine CCL3 production. Glia. 2015;63(1):51–65. https://api.istex.fr/ark:/67375/WNG-F56FH48S-N/fulltext.pdf. doi: 10.1002/glia.22732.

108. Nakagawa S, Izumi Y, Takada-Takatori Y, Akaike A, Kume T. Increased CCL6 expression in astrocytes and neuronal protection from neuron–astrocyte interactions. Biochemical and Biophysical Research Communications. 2019;519(4):777–782. https://dx.doi.org/10.1016/j.bbrc.2019.09.030. doi: 10.1016/j.bbrc.2019.09.030.

109. Feng X, Ji Y, Zhang C, Jin T, Li J, Guo J. CCL6 promotes M2 polarization and inhibits macrophage autophagy by activating PI3-kinase/akt signalling pathway during skin wound healing. Experimental dermatology. 2023;32(4):403–412. https://onlinelibrary.wiley.com/doi/abs/10.1111/exd.14718. doi: 10.1111/exd.14718.

110. Fukuda M, Mikoshiba K. Doc2gamma, a third isoform of double C2 protein, lacking calcium-dependent phospholipid binding activity. Biochemical and biophysical research communications. 2000;276(2):626–632. https://www.ncbi.nlm.nih.gov/pubmed/11027523.

111. Friedrich R, Yeheskel A, Ashery U. DOC2B, C2 domains, and calcium: A tale of intricate interactions. Mol Neurobiol. 2010;41(1):42–51. https://link.springer.com/article/10.1007/s12035-009-8094-8. doi: 10.1007/s12035-009-8094-8.

112. Francis PT. Glutamatergic systems in alzheimer’s disease. Int J Geriatr Psychiatry. 2003;18:15.

113. Nagatsu T, Nakashima A, Watanabe H, Ito S, Wakamatsu K. Neuromelanin in parkinson’s disease: Tyrosine hydroxylase and tyrosinase. International Journal of Molecular Sciences. 2022;23(8):4176. https://www.ncbi.nlm.nih.gov/pubmed/35456994. doi: 10.3390/ijms23084176.

114. Aliev G, Shahida K, Gan SH, et al. Alzheimer disease and type 2 diabetes mellitus: The link to tyrosine hydroxylase and probable nutritional strategies. CNS & neurological disorders drug targets. 2014;13(3):467–477. http://www.eurekaselect.com/openurl/content.php?genre=article&issn=1871-5273&volume=13&issue=3&spage=467. doi: 10.2174/18715273113126660153.

115. Li J, Wang X, Chen L, et al. TMEM158 promotes the proliferation and migration of glioma cells via STAT3 signaling in glioblastomas. Cancer gene therapy. 2022;29(8-9):1117–1129. https://www.ncbi.nlm.nih.gov/pubmed/34992215. doi: 10.1038/s41417-021-00414-5.

116. Tong J, Li H, Hu Y, Zhao Z, Li M. TMEM158 regulates the canonical and non-canonical pathways of TGF-β to mediate EMT in triple-negative breast cancer. Journal of Cancer. 2022;13(8):2694–2704. https://search.proquest.com/docview/2678427046. doi: 10.7150/jca.65822.

117. Fu Y, Yao N, Ding D, et al. TMEM158 promotes pancreatic cancer aggressiveness by activation of TGFβ1 and PI3K/AKT signaling pathway. Journal of cellular physiology. 2020;235(3):2761–2775. https://onlinelibrary.wiley.com/doi/abs/10.1002%2Fjcp.29181. doi: 10.1002/jcp.29181.

118. Fukuchi K, Hart M, Li L. Alzheimer’s disease and heparan sulfate proteoglycan. Frontiers in bioscience. 1998;3:327. https://www.ncbi.nlm.nih.gov/pubmed/9490646. doi: 10.2741/A277.

119. Snow AD, Cummings JA, Lake T. The unifying hypothesis of alzheimer’s disease: Heparan sulfate proteoglycans/glycosaminoglycans are key as first hypothesized over 30 years ago. Frontiers in aging neuroscience. 2021;13:710683. https://search.proquest.com/docview/2578914403. doi: 10.3389/fnagi.2021.710683.

120. Kolev MV, Ruseva MM, Harris CL, Morgan BP, Donev RM. Implication of complement system and its regulators in alzheimers disease. Current Neuropharmacology. 2009;7(1):1–8. http://www.eurekaselect.com/openurl/content.php?genre=article&issn=1570159X&volume=7&issue=1&spage=1. doi: 10.2174/157015909787602805.

121. Zhao L, Berra HH, Duan W, et al. Beneficial effects of hematopoietic growth factor therapy in chronic ischemic stroke in rats. Stroke. 2007;38(10):2804–2811. http://stroke.ahajournals.org/cgi/content/abstract/38/10/2804. doi: 10.1161/STROKEAHA.107.486217.

122. Zhao L, Singhal S, Duan W, Mehta J, Kessler JA. Brain repair by hematopoietic growth factors in a rat model of stroke. Stroke. 2007;38(9):2584–2591. http://stroke.ahajournals.org/cgi/content/abstract/38/9/2584. doi: 10.1161/STROKEAHA.106.476457.

123. Cui L, Murikinati SR, Wang D, Zhang X, Duan W, Zhao L. Reestablishing neuronal networks in the aged brain by stem cell factor and granulocyte-colony stimulating factor in a mouse model of chronic stroke. PloS one. 2013;8(6):e64684. https://www.ncbi.nlm.nih.gov/pubmed/23750212. doi: 10.1371/journal.pone.0064684.

124. Piao C, Gonzalez-Toledo ME, Xue Y, et al. The role of stem cell factor and granulocyte-colony stimulating factor in brain repair during chronic stroke. Journal of Cerebral Blood Flow & Metabolism. 2009;29(4):759–770. https://dx.doi.org/10.1038/jcbfm.2008.168. doi: 10.1038/jcbfm.2008.168.

125. Facon T, Harousseau J, Maloisel F, et al. Stem cell factor in combination with filgrastim after chemotherapy improves peripheral blood progenitor cell yield and reduces apheresis requirements in multiple myeloma patients: A randomized, controlled trial. Blood. 1999;94(4):1218–1225. https://dx.doi.org/10.1182/blood.V94.4.1218. doi: 10.1182/blood.V94.4.1218.

126. Stiff P, Gingrich R, Bensinger W, et al. A randomized phase 2 study of PBPC mobilization by stem cell factor and filgrastim in heavily pretreated patients with hodgkin’s disease or non-hodgkin’s lymphoma. Bone marrow transplantation. 2000;26(5):471–481. https://www.ncbi.nlm.nih.gov/pubmed/11019835. doi: 10.1038/sj.bmt.1702531.

127. 2024 alzheimer’s disease facts and figuresAlzheimer’s & dementia. 2024;20(5):3708–3821. https://onlinelibrary.wiley.com/doi/abs/10.1002%2Falz.13809. doi: 10.1002/alz.13809.

128. Duarte RF, Frank DA. The synergy between stem cell factor (SCF) and granulocyte colony-stimulating factor (G-CSF): Molecular basis and clinical relevance. Leukemia & Lymphoma. 2002;43(6):1179–1187. https://www.tandfonline.com/doi/abs/10.1080/10428190290026231. doi: 10.1080/10428190290026231.

129. To LB, Bashford J, Rawling T, et al. Successful mobilization of peripheral blood stem cells after addition of ancestim (stem cell factor) in patients who had failed a prior mobilization with filgrastim (granulocyte colony-stimulating factor) alone or with chemotherapy plus filgrastim. Bone marrow transplantation. 2003;31(5):371–378. https://dx.doi.org/10.1038/sj.bmt.1703860. doi: 10.1038/sj.bmt.1703860.

