## Supplementary material for "Transcriptomic analysis reveals new reparative mechanisms of SCF+G-CSF-reduced neuropathology in aged APP/PS1 mice": Suppemental figures and table

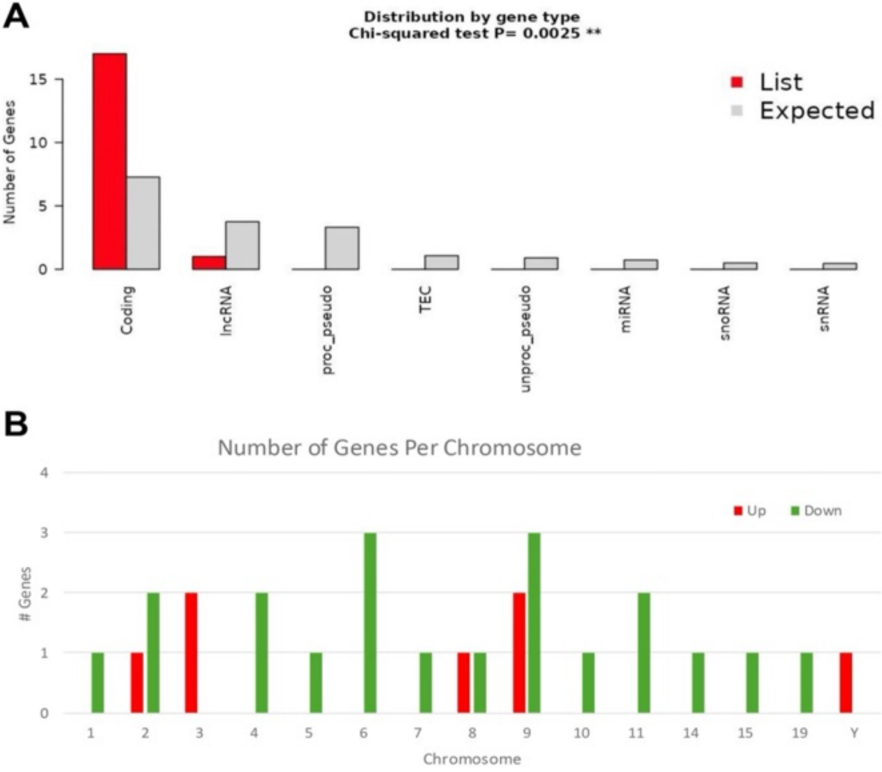


**Figure S1**. Chromosomal analysis of SCF+G-CSF-induced differentially expressed genes (DEGs) using ShinyGO 0.77. (**A**) SCF+G-CSF-induced DEG analysis reveals a significantly higher proportion of protein-coding genes and a lower proportion of long non-coding RNA (lncRNA) genes than expected by chance. (**B**) Chromosomal mapping of DEGs shows that SCF+G-CSF treatment in aged APP/PS1 mice results in more downregulated than upregulated genes across the genome.

**A** **B**

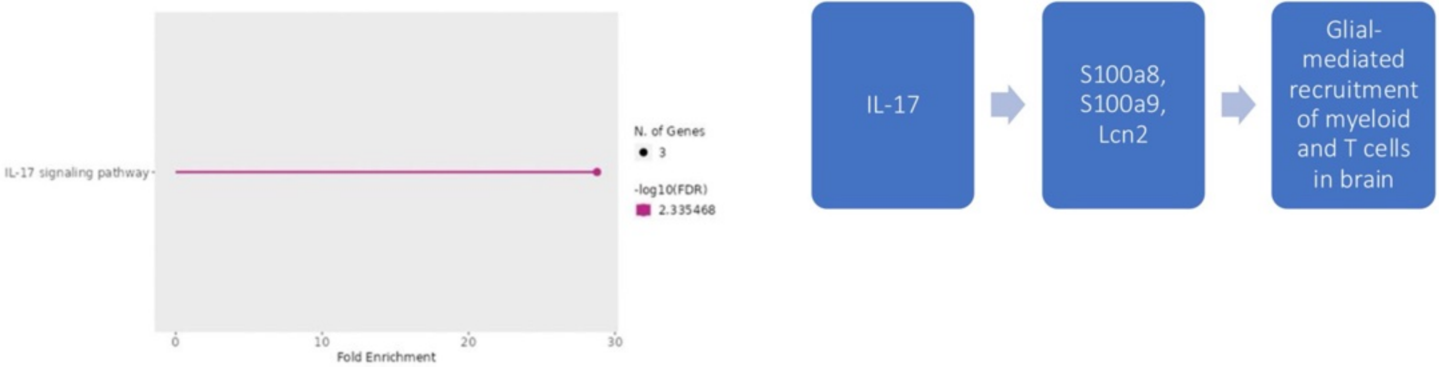


| Enrichment FDR | nGenes | Pathway Genes | Fold Enrichment | Pathway |
|---|---|---|---|---|
| 4.5E-03 | 3 | 91 | 29.1 | IL-17 signaling pathway |

**Figure S2**. SCF+G-CSF-induced enrichment of the IL-17 signaling pathway stimulates S100a8, S100a9, and Lcn2 to promote immune cell infiltration into the brains of aged APP/PS1 mice. (**A**) ShinyGO 0.77 analysis reveals a 29.1-fold enrichment of the IL-17 signaling pathway following SCF+G-CSF treatment, with 3 of 91 pathway-associated genes (S100a8, S100a9, and Lcn2) identified in the dataset. (**B**) This IL-17 pathway enrichment is further supported by STRING v12.0 analysis (see Figure 4). We propose that SCF+G-CSF-induced activation of S100a8, S100a9, and Lcn2 via IL-17 signaling facilitates infiltration of innate and adaptive immune cells into the brain, a process that may contribute to the reduction of AD-related neuropathology.

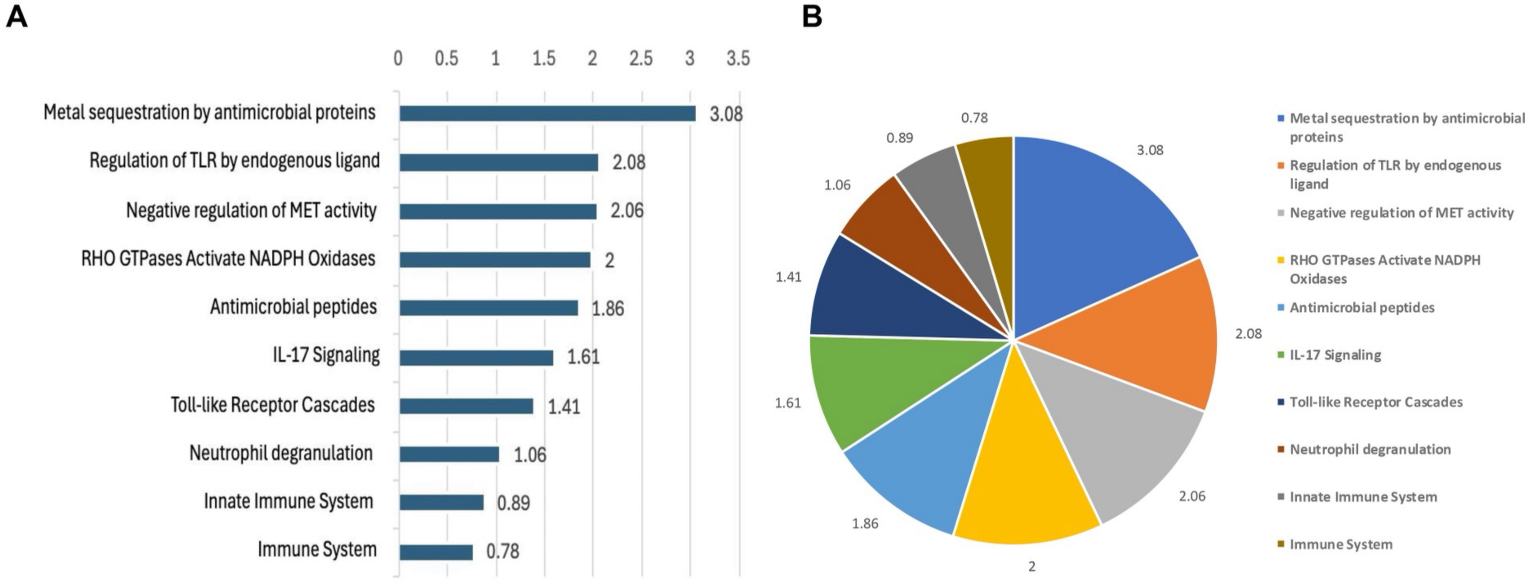


**Figure S3**. SCF+G-CSF-treatment promotes metal sequestration and immune system functions in aged APP/PS1 mice. STRING v12.0 pathway enrichment analysis ranked by strength in SCF+G-CSF-treated aged APP/PS1 mouse brains. (**A**) SCF+G-CSF treatment promotes enrichment of pathways related to metal sequestration and various immune system functions in the brains of aged APP/PS1 mice. (**B**) Pie chart representation of pathway enrichment, organized by strength, highlights the proportional contribution of each functional category.

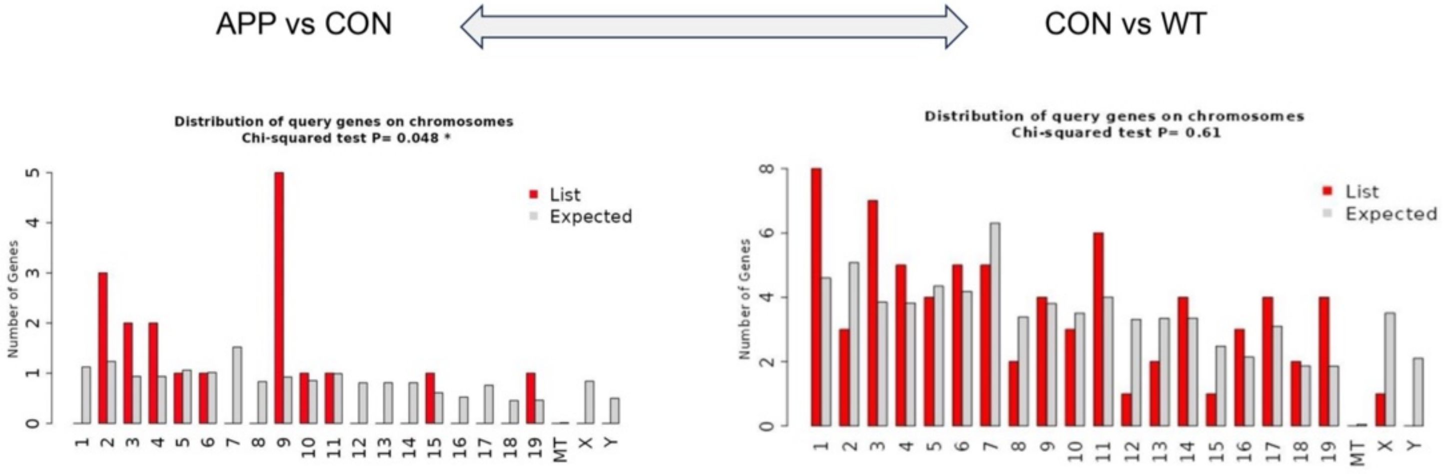


**Figure S4.** Chromosomal distribution of gene expression in aged APP/PS1 mice reveals loss of transcriptional control without SCF+G-CSF treatment. Chromosomal mapping of differentially expressed genes (DEGs) using ShinyGO 0.77 shows widespread gene expression across nearly all chromosomes in aged APP/PS1 mice treated with vehicle. In contrast, SCF+G-CSF treatment markedly reduces or suppresses gene expression across multiple chromosomes, indicating a partial restoration of transcriptional control disrupted in the AD state. APP: aged APP/PS1 mice treated with SCF+G-CSF; CON: aged APP/PS1 mice injected with vehicle; WT: age-matched wild-type mice.

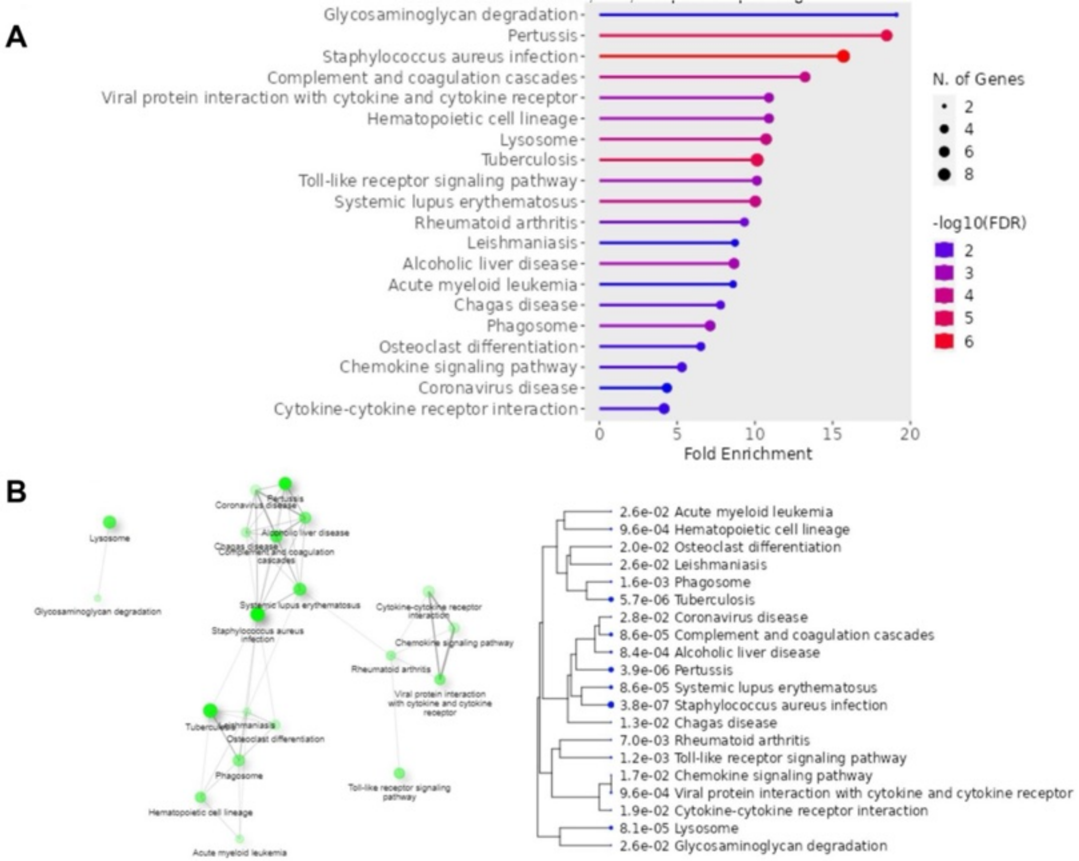


**Figure S5**. ShinyGO 0.77 functional enrichment analysis reveals glycosaminoglycan degradation as the top-enriched pathway in aged APP/PS1 control mice. Functional enrichment was performed on 75 of the 89 differentially expressed genes (DEGs) identified in aged APP/PS1 control mice compared to WT mice. (**A**) Pathways are ranked by fold enrichment, with glycosaminoglycan degradation showing the highest enrichment. (**B**) Interaction network highlights functional connections among enriched genes, including those linked to bacterial, viral, and parasitic infection-related pathways.

**Table S1.** Chromosomal localization of SCF+G-CSF-induced DEGs

| ID count: 27 | Gene Symbol | Description | Chro... | Start | Stop |
|---|---|---|---|---|---|
| 1418188_a_at | Malat1 | metastasis associated lun... | chr11 | 101246423 | 101248246 |
| 1418722_at | Ngp | neutrophilic granule prot... | chr9 | 110419845 | 110423015 |
| 1419394_s_at | S100a8 | S100 calcium binding pro... | chr3 | 90669070 | 90670034 |
| 1419691_at | Camp | cathelicidin antimicrobial... | chr9 | 109847378 | 109849617 |
| 1427747_a_at | Lcn2 | lipocalin 2 | chr2 | 32384636 | 32387739 |
| 1427820_at | | | chrY | 90785641 | 90816464 |
| 1436746_at | Wnk1 | WNK lysine deficient prot... | chr6 | 119954046 | 120037398 |
| 1437082_at | Akap9 | A kinase (PRKA) anchor p... | chr5 | 3928185 | 3960475 |
| 1438403_s_at | Malat1 | metastasis associated lun... | chr19 | 5802111 | 5802678 |
| 1448756_at | S100a9 | S100 calcium binding pro... | chr3 | 90692631 | 90695298 |
| 1450009_at | Ltf | lactotransferrin | chr9 | 111019302 | 111042766 |
| 1450906_at | Plxnc1 | plexin C1 | chr10 | 94790861 | 94944578 |
| 1454696_at | Gnb1 | guanine nucleotide bindi... | chr4 | 155557943 | 155559269 |
| 1455886_at | Cbl | Casitas B-lineage lympho... | chr9 | 44149247 | 44150358 |
| 1430309_at | Nipbl | Nipped-B homolog (Dros... | chr15 | 8359158 | 8444463 |
| 1431060_at | Peli1 | pellino 1 | chr11 | 21091275 | 21143005 |
| 1437311_at | Snhg11 | small nucleolar RNA host... | chr2 | 158375660 | 158382080 |
| 1439566_at | Gprin3 | GPRIN family member 3 | chr6 | 59352450 | 59353513 |
| 1440034_at | Stam2 | signal transducing adapt... | chr2 | 52699499 | 52700173 |
| 1440161_at | Mmp16 | matrix metallopeptidase 16 | chr4 | 18118819 | 18119145 |
| 1440290_at | | | chr6 | 128187831 | 128188377 |
| 1440314_at | | | chr1 | 84747717 | 84748382 |
| 1440464_at | | | chr8 | 4309913 | 4310376 |
| 1441606_at | | | chr14 | 38954475 | 38955114 |
| 1442029_at | Kcnq1ot1 | KCNQ1 overlapping trans... | chr7 | 143242337 | 143243585 |
| 1452722_a_at | Cul5 | cullin 5 | chr9 | 53614581 | 53667488 |
| 1457577_at | | | chr8 | 8642762 | 8643346 |
